# CARPOOL: A library-based platform to rapidly identify next generation chimeric antigen receptors

**DOI:** 10.1101/2021.07.09.450900

**Authors:** Taeyoon Kyung, Khloe S. Gordon, Caleb R. Perez, Patrick V. Holec, Azucena Ramos, Angela Q. Zhang, Yunpeng Liu, Catherine Koch, Alina Starchenko, Brian Joughin, Douglas A. Lauffenburger, Darrell J. Irvine, Michael T. Hemann, Michael E. Birnbaum

**Affiliations:** Koch Institute for Integrative Cancer Research, Cambridge, MA, USA; Department of Biological Engineering, Massachusetts Institute of Technology, Cambridge, MA, USA; Singapore-MIT Alliance for Research and Technology Centre, Singapore 138602, Singapore; Department of Biology, Massachusetts Institute of Technology, Cambridge, MA, USA; Department of Health, Science, and Technology, Massachusetts Institute of Technology, Cambridge, MA, USA; Ragon Institute of MIT, MGH, and Harvard, Cambridge, MA, USA

## Abstract

CD19-targeted CAR therapies have successfully treated B cell leukemias and lymphomas, but many responders later relapse or experience toxicities. CAR intracellular domains (ICDs) are key to converting antigen recognition into anti-tumor effector functions. Despite the many possible immune signaling domain combinations that could be included in CARs, almost all CARs currently rely upon CD3**ζ**, CD28, and/or 4-1BB signaling. To explore the signaling potential of CAR ICDs, we generated a library of 700,000 CD19 CAR molecules with diverse signaling domains and developed a high throughput screening platform to enable optimization of CAR signaling for anti-tumor functions. Our strategy identifies CARs with novel signaling domain combinations that elicit distinct T cell behaviors from a clinically available CAR, including enhanced proliferation and persistence, lower exhaustion, potent cytotoxicity in an *in vitro* tumor rechallenge condition, and comparable tumor control *in vivo*. This approach is readily adaptable to numerous disease models, cell types, and selection conditions, making it a promising tool for rapidly improving adoptive cell therapies and expanding their utility to new disease indications.

## Main Text

Cancer immunotherapies that reinvigorate or reprogram anti-tumor T cell responses have been transformative in the treatment of a broad range of malignancies.^1^ Chimeric antigen receptors (CARs) engineer T cells to leverage these mechanisms by linking intracellular immunostimulatory signaling domains to extracellular recognition domains typically derived from antibody single chain variable fragments (scFv). Upon stable introduction into T cells, the CAR redirects effector functions toward a clinically relevant target antigen. This has been most widely utilized in the context of B cell malignancies, where a majority of patients show complete responses within the first few months.^2^ However, many patients later relapse, and most patients experience cytokine release syndrome or neurological impairment.^2^ Further, CAR-T treatments have yet to meaningfully translate to solid tumors, which pose distinct immunological challenges.

Retrospective clinical studies have established which CAR T cell phenotypes are beneficial for long-term efficacy and safety.^2,3^ These phenotypes can be induced via the composition of the CAR signaling domains. While intracellular domains (ICDs) from CD3**ζ** in combination with CD28 and/or 4-1BB are most commonly used in current CARs,^3,4^ several groups have shown that incorporating novel signaling components can enhance persistence, proliferation, cytotoxicity, resistance to exhaustion, memory formation, and *in vivo* survival benefit.^5–7^ Additionally, shortening the distance between the CD3**ζ** immunoreceptor tyrosine activation motifs (ITAMs) and the membrane can enhance their function, and it has been demonstrated that a CAR containing a single ITAM produced superior persistence and *in vivo* tumor control than the canonical sequence.^8^ The effects of distinct signaling domains can also synergize when arranged in the optimal spatial configuration.^9–11^ While there is a steadily expanding compendium of CARs utilizing novel signaling domains to confer functions, these compositions undersample all possible signaling domain combinations that could be beneficial in CAR constructs, possibly impeding efficacy and translation to other diseases.

The relative scarcity of tested signaling domain combinations is a result of the time and effort needed to individually design and test new CARs. We hypothesized that a more systematic optimization of CAR signaling domains has the potential to produce novel CAR-T cell behaviors that could elicit safer, more effective therapies. To this end, we created a 700,000-member CAR library with diversified ICDs and coupled it with a selection strategy that enables enrichment of CAR-T cells exhibiting desirable anti-tumor functions (which, together, we term CARPOOL). We identified several novel CAR ICD combinations, and show that one in particular shows superior activity relative to a conventional 19BB**ζ** CAR (BB**ζ**). Taken together, this evidence suggests that CARPOOL can rapidly optimize CAR ICDs to enhance therapeutic function and poses a promising strategy to address current challenges faced by CAR-T therapies.^2^

## Results

### CARPOOL streamlines selection of novel high-performance CARs

In order to design a CAR library, we identified 89 signaling domains derived from different immune cell types and functional families (**Supplementary Table 1**), which were then incorporated at random into each of the three intracellular positions in a 3rd generation CD19 CAR construct. Each CAR was cloned into a lentiviral transfer vector along with a randomized 18-nucleotide barcode sequence in the 3’UTR, yielding a theoretical diversity of 700,000 uniquely barcoded signaling domain combinations (**Fig. 1a**). We then generated lentiviruses from the CAR plasmid library, and transduced 1×10^8^ Jurkat T cells at a multiplicity of infection (MOI) of 0.5 to favor incorporation of a single CAR-encoding transgene per cell. This produced 3×10^7^ CAR library transduced cells, as confirmed by epitope tag staining of the CAR extracellular region, achieving approximately 40-fold coverage of our theoretical library size.

**Figure 1.**
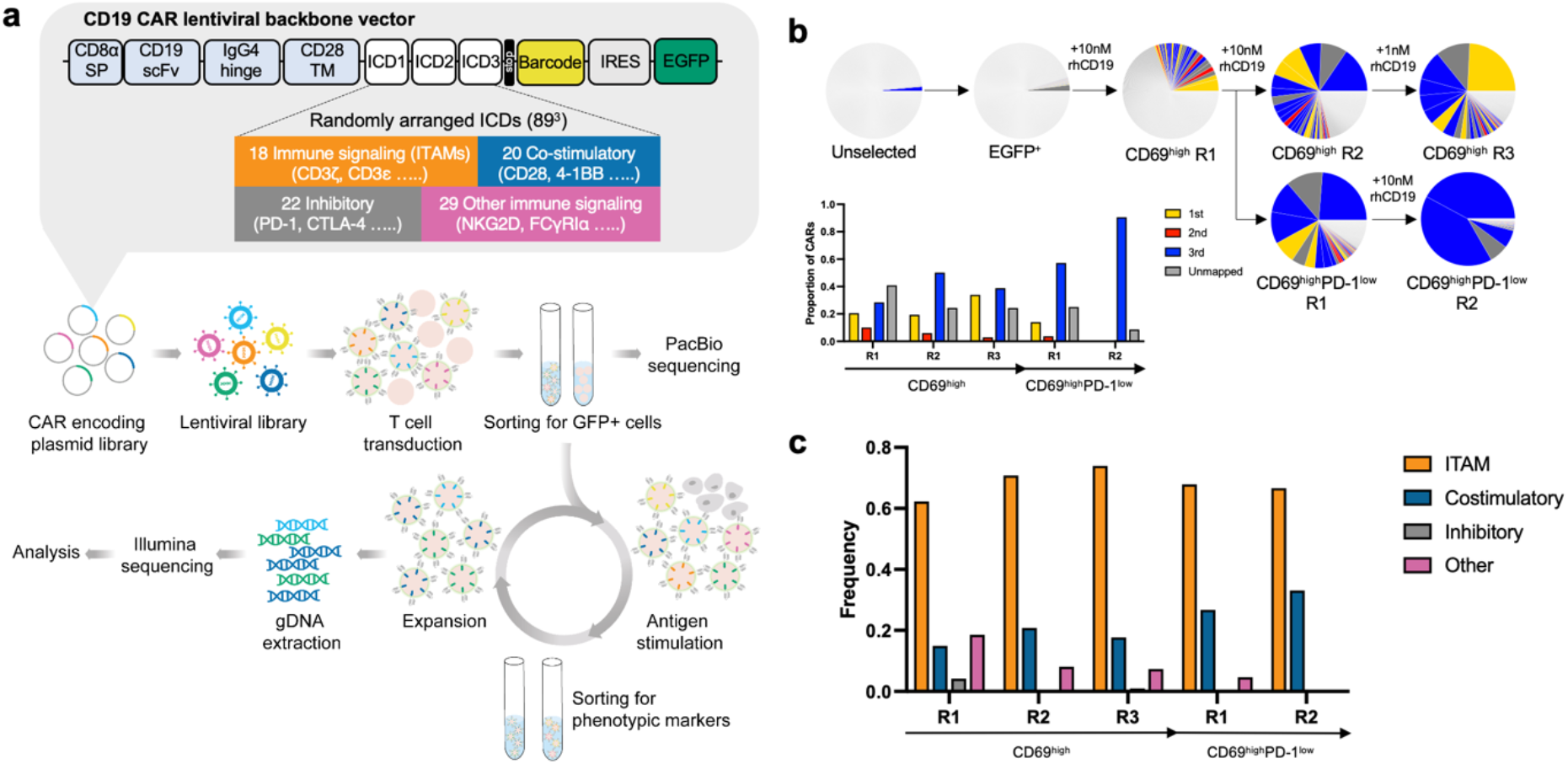
Selecting for CD69^high^ expression enriches CARs encompassing ITAM-signaling ICDs. (**a**) Schematics describing the CARPOOL system. CARPOOL utilizes a signaling diversified library of CD19-specific chimeric antigen receptors containing 1-3 ICDs and combines cell sorting and next generation sequencing to select for function in human T cells. All CARs were bi-cistronically expressed with EGFP via an IRES sequence. (**b**) Frequency of top 30 CAR clones throughout rounds of selection in Jurkat T cells for CD69^high^ and CD69^high^PD-1^low^ expression following stimulation with 10 nM rhCD19 with the proportion of 1st, 2nd, and 3rd generation CARs identified. (**c**) Frequency of each family of ICDs throughout rounds of selection.

We next FACS sorted for cells EGFP expression, which was bi-cistronically expressed with each CAR construct. For the first round of selection, we stimulated the EGFP-enriched T cell pool with 10 nM of soluble human CD19 recombinant protein (a.a. 1-270) (rhCD19) and subsequently sorted the stimulated EGFP^+^ cells that expressed the highest levels of CD69, a canonical T cell activation maker (CD69^high^).^12,13^ We then split this selected population in two in order to conduct two more rounds of selections in serial for either CD69^high^ expression to further enrich for robustly activatable CARs, or for CD69^high^PD-1^low^ expression to potentially identify ICD combinations that render T cells less susceptible to exhaustion. The proportion of CD69^+^ cells increased substantially throughout rounds of both CD69^high^ and CD69^high^PD-1^low^ selections, indicating an enrichment for functional CAR-T cells: 72% and 90% of total cell populations were EGFP^+^CD69^+^ cells after the last round of CD69^high^ and CD69^high^PD-1^low^ selections, respectively (**Supplementary Fig. 1**).

Next, we performed next-generation sequencing (NGS) to track the prevalence of enriched barcodes in selected cells from each round of selection. We observed a dramatic reduction in clonal diversity following each round of both CD69^high^ and CD69^high^PD-1^low^ selections, where the top 25 most enriched barcodes represented 89% and 99% of all clones from the last rounds of CD69^high^ and CD69^high^PD-1^low^ selections, respectively (**Supplementary Fig. 2**). Given the limited read length capacity of Illumina sequencing, we used PacBio SMRT long-read sequencing of amplicons derived from CARPOOL transduced Jurkats that encompass both the CAR ICDs and barcode region in order to build a lookup table to link the enriched barcodes to the ICD combinations. Of note, only ~70% of barcodes in the top 30 most frequent CARs from all selections were identified in the PacBio data, potentially due to read depth limitations in the SMRT sequencing. Additionally, not all identified CARs encompassed 3 ICDs as intended, likely due to infidelities in the library assembly step (**Fig. 1b**).

### Enriched CARs encode ITAM-containing ICDs with unique combinations of costimulatory ICDs

We quantified the average frequency of each ICD at each selection step to determine which classes of ICDs were most enriched. Our result verified that ITAM-containing ICDs, which are the canonical activation motifs for antigen receptor signaling,^14,15^ were more prevalent than other classes of ICDs, especially in later rounds of CD69-based selection, consistent with a requirement for ITAMs for robust CAR activation (**Fig. 1c**).

Analysis of our sequencing data revealed enrichment of novel signaling domains that have not been vetted for function in a CAR, alongside commonly used ICDs such as CD28 and 41BB and ICDs that have been more recently described as functional.^7,16–20^ However, it is notable that we identified a BB**ζ** CAR clone in our library that was less enriched compared to our top clones, especially in the last rounds of both CD69^high^ and CD69^high^PD-1^low^ selections, indicating that these novel clones possess competitive advantages over the BB**ζ** CAR in these selection conditions (**Fig. 2a**). In order to visualize and track the extent of enrichment for each ICD, we generated heatmaps for each round of selection that depict the frequency of each ICD at each position (**Fig. 2b** and **Supplementary Fig. 3**). Furthermore, our analysis suggested that previously uncharacterized combinations of co-stimulatory and ITAM-containing ICDs, such as FcεR1y, 2B4, and CD3ε ITAM or CD79b, DAP12, and CD40, were overrepresented following selection relative to more commonly used signaling domain combinations utilizing 4-1BB, CD28, and CD3**ζ** (**Fig. 2a**). This exemplifies the utility of CARPOOL in revealing useful signaling domain combinations in a rapid, streamlined process. Notably, our enriched data identified ICDs that are contained in FDA approved 2nd generation CARs (CD28, 4-1BB, and CD3**ζ**), as well as domains in clinical trials or under consideration for novel 2nd and 3rd generation CARs (**Fig. 2b** and Supplementary **Fig. 3**).

**Figure 2.**
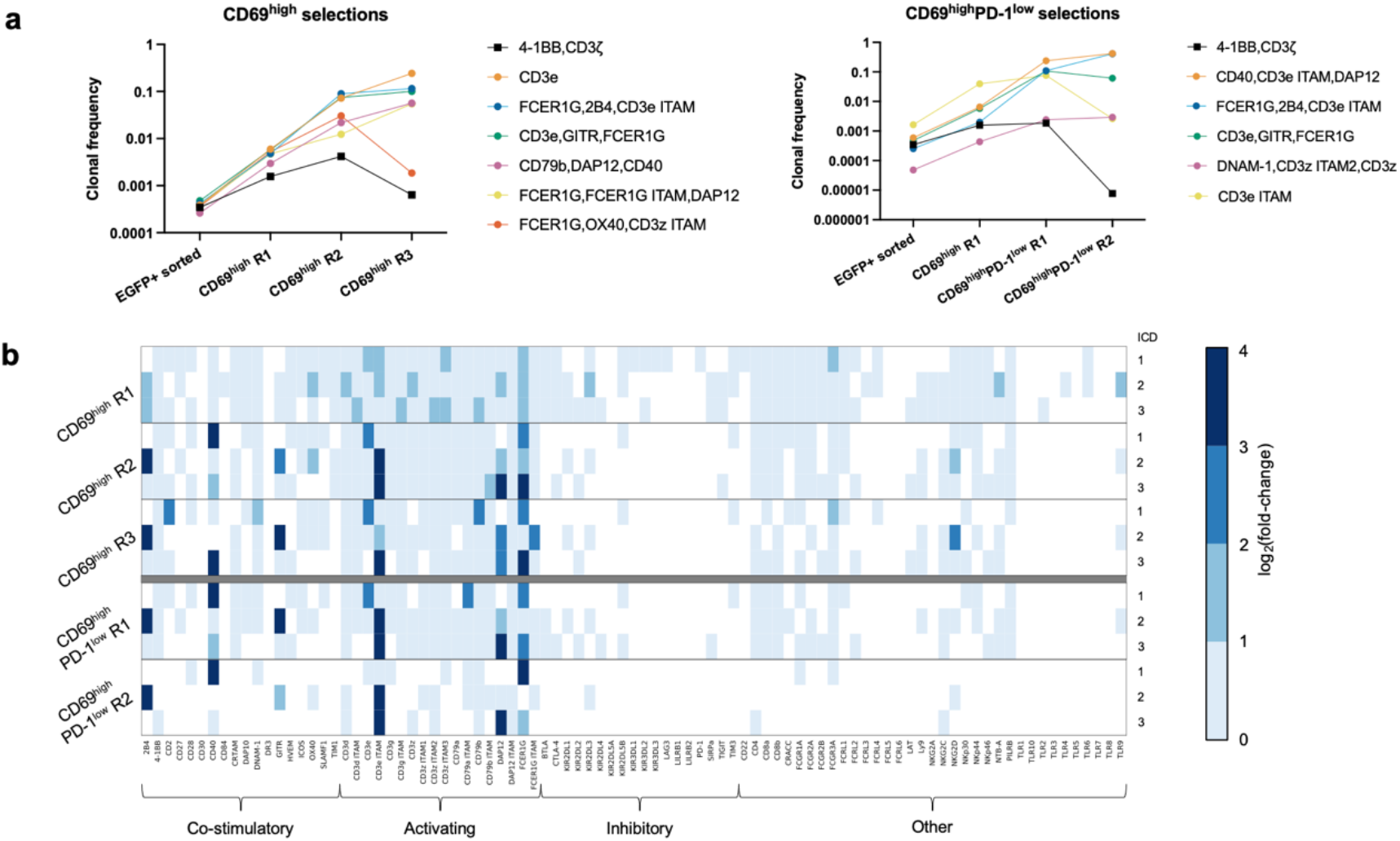
CARPOOL selections reveal novel functional signaling domain configurations. (**a**) Clonal frequency of enriched CARs that were identified throughout rounds of CD69^high^ and CD69^high^PD-1^low^ selections compared to a BB**ζ** CAR. (**b**) Heatmap representing log_2_(fold-change) of individual ICDs found in 3rd generation CARs throughout rounds of selection, oriented by ICD position within the CAR intracellular domain relative to the plasma membrane.

### CAR variants from library selections show distinct functional phenotypes compared to those of a BBζ CAR in primary T cells

In order to compare the functional characteristics of our novel CARs to those of a BB**ζ** CAR, we vetted 3 highly enriched and distinct CARs in Jurkat T cells. While we detected comparable surface expression levels and antigen sensitivity upon assessing both CAR internalization and CD69 up-regulation relative to BB**ζ** (**Supplementary Fig. 4a–d**), we noted that the CD69 and PD-1 expression levels of all three variants in the absence of antigen engagement were significantly lower than those of the BB**ζ** CAR (**Supplementary Fig. 4e–f**). This indicates that they induce less tonic signaling in Jurkats—a feature that has been reported to lead to rapid CAR T cell dysfunction marked by deficient IL-2 and IFN-γ production.^21,22^ Upon antigen stimulation, we noted that two of the enriched CARs, Var1 and Var3, showed robust CD69 upregulation while Var2 consistently produced lower CD69 upregulation (**Supplementary Fig. 4d**).

Thus, we chose to further characterize the two more active hits from our library selections: Var1 (containing ICDs from CD40, CD3ε ITAM, and DAP12), which was highly enriched in both CD69^high^ and CD69^high^PD-1^low^ selections, and Var3 (containing ICDs from FcεR1γ, OX40, and CD3**ζ** ITAM3), which was enriched in the CD69^high^ selection but not in the CD69^high^PD-1^low^ selection (**Fig. 3a**). These were chosen not only for their unique signaling compositions and relative enrichment, but also to explore whether a PD-1^low^ selection criteria enabled identification of CARs with reduced susceptibility to T cell exhaustion.

**Figure 3.**
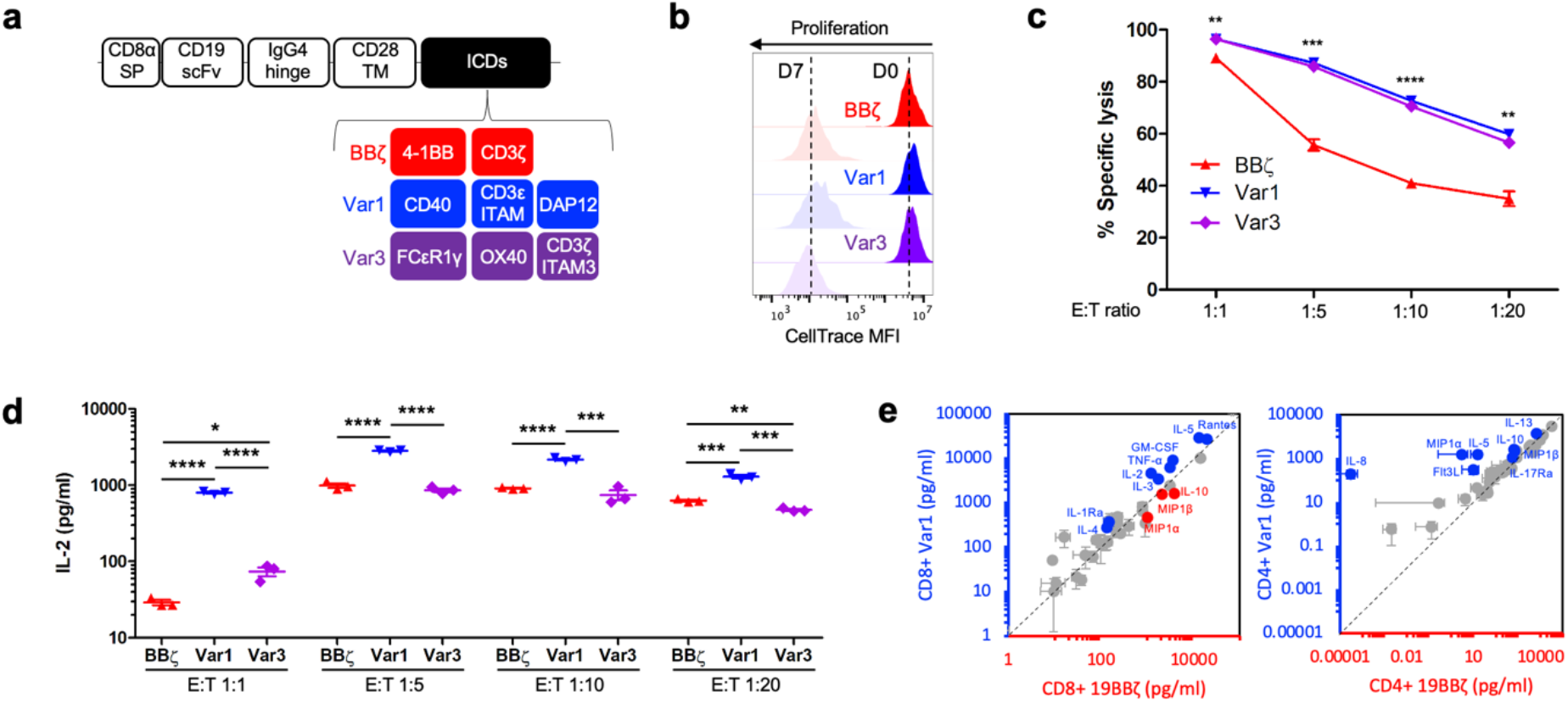
CAR variants show enhanced cytotoxicity and cytokine secretion in response to antigen stimulation. (**a**) Design of CD19 targeted CAR candidates. (**b**) Proliferation of human primary CD8^+^ CAR T cells that were stained with CellTrace dye on day 0 prior to co-culture with CD19^+^ NALM6 cells at an E:T ratio of 1:1 (n = 3 technical replicates representative of 2 biological replicates). Extent of proliferation was assessed by degree of dye dilution and measured by FACS. (**c**) Cytotoxicity and (**d**) IL-2 secretion from human primary CD8^+^ CAR T cells following 24 hour co-culture with FLuc^+^ CD19^+^ NALM6 cells at varying E:T ratios (n = 3 technical replicates representative of 2 biological replicates). Remaining NALM6 cells were quantified by measuring bioluminescent activity, while IL-2 levels were measured via ELISA. *P* values in panel (**c**) are 0.001820, 0.000140, <0.000001, and 0.001293 for Var1 vs. BB**ζ** and 0.001860, 0.000197, 0.000007, and 0.002098 for Var3 vs. BB**ζ** in increasing order of E:T ratio, as determined using a two-tailed unpaired student’s t-test with Benjamini-Hochberg correction (df = 4 for all). Data and error bars shown are means ± s.e.m. *P* values in panel (**d**) are <0.0001 for Var1 vs. BB**ζ**, 0.0113 for BB**ζ** vs. Var3, and < 0.0001 for Var3 vs. BB**ζ**, and 0.0008 for Var1 vs. Var3 at an E:T of 1:1 ratio (df = 4). *P* values are < 0.0001 for Var1 vs. BB**ζ** and <0.0001 Var1 vs. Var3 at an E:T 1:5 of ratio (df = 4). *P* values are < 0.0001 for Var1 vs. BB**ζ** and 0.0004 for Var1 vs. Var3 at an E:T of 1:10 ratio (df = 4). *P* values are 0.0007 for Var1 vs. BB**ζ**, 0.0024 for Var3 vs. BB**ζ**, and 0.0003 for Var1 vs. Var3 at E:T 1:20 ratio (df = 4). *P* values for panel (**d**) were determined using a two-tailed unpaired student’s t-test. Data shown are individual points along with means ± s.e.m. (**e**) Polyfunctional cytokine and chemokine secretion response following 24 hour co-culture of human primary CD8^+^ or CD4^+^ CAR T cells (n = 3 technical replicates). Concentrations were quantified for 41 analytes. Cytokines linked to anti-tumor response that were produced at significantly higher levels by the novel CARs relative to 19BB**ζ** are highlighted in blue, while those that were secreted at lower levels are highlighted in red.

Given that the range of functional phenotypes following CAR activation is limited in Jurkats, we next characterized the function of each CAR variant in human primary T cells. While we did not observe meaningful differences between the CAR variants and BB**ζ** in the absence of antigen (**Supplementary Fig. 5**), we speculated that differences in phenotype between Var1 and Var3 from BB**ζ** may be driven by antigen-induced CAR signaling. CD8^+^ T cells expressing Var1, Var3, and BB**ζ** were co-cultured with CD19-expressing NALM6 cells. While we found no difference in T cell proliferation at day 7 (**Fig. 3b**), there were significant differences in killing capacity following 24 hour co-culture with NALM6 cells, with Var1 and Var3 exhibiting increased cytotoxicity compared to BB**ζ** across all E:T ratios, and especially at high tumor burden (**Fig. 3c**). This trend was matched with elevated levels of IL-2 secretion by Var1 at all E:T ratios (**Fig. 3d**), but with no observed differences in IFN-γ secretion (**Supplementary Fig. 6**).

To assess whether there were further notable differences in cytokine secretion between CARs, we conducted a 41-plex Luminex assay using supernatants collected after co-culture of either CD4^+^ or CD8^+^ CAR-T cells with NALM6 cells at an E:T ratio of 1:1. We detected increased secretion of cytokines associated with anti-tumor effects by both Var1 and Var3 relative to BB**ζ**, such as IL-2, TNF-a, and GM-CSF, in CD8^+^ T cells (**Fig. 3e and Supplementary Fig. 7**). In CD4^+^ T cells, we found elevated levels of chemokines associated with attracting immune cells from both the innate and adaptive immune system compared to BB**ζ**, including MIP1α, MIP1β, and Flt3L in the case of Var1 and MIP1α and IP-10 in the case of Var3, indicating that modification of CAR signaling can drastically alter their cytokine secretion profiles in T cells.^23–27^

### Var1 CAR-T cells show enhanced persistence and tumor control in a long-term tumor rechallenge condition

In order to assess how the enhanced killing and anti-tumor cytokine secretion displayed by CARPOOL-enriched CAR variants would affect function over an extended period of exposure to high tumor burden, we designed a rechallenge assay in which we repeatedly added NALM6 cells every 2-3 days at an E:T ratio of 1:10 to a 1:1 mixture of CD4^+^ and CD8^+^ T cells expressing each CAR (**Fig. 4a**). Under these conditions, we found that Var1-expressing CD4^+^ and CD8^+^ T cells demonstrated considerably more persistent proliferative activity at later time points compared to both Var3- and BB**ζ**-expressing cells (**Fig. 4b**). This expansion pattern directly correlated with tumor control, with Var1 showing superior tumor control at late timepoints (**Fig. 4c**). Var1-expressing cells also showed delayed kinetics of differentiation from T cell memory to effector phenotypes along with significantly reduced exhaustion marker expression levels by day 22 (**Fig. 4d–e and Supplementary Fig. 8**). Taken together, these results imply that Var1-expressing T cells are less susceptible to developing an exhausted T cell phenotype upon repeated challenge with high tumor burden relative to Var3 and BB**ζ** CAR-T cells.

**Figure 4.**
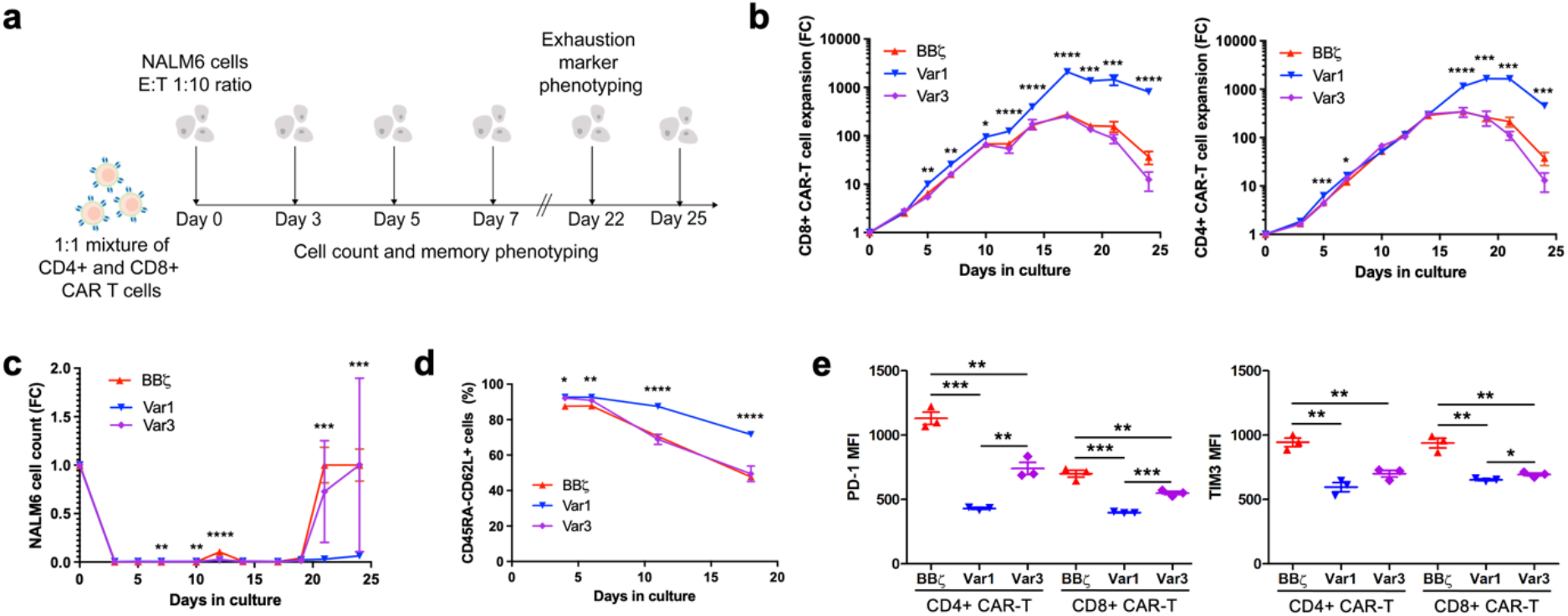
Var1 demonstrates enhanced persistence and anti-tumor function with lower exhaustion following long term tumor challenge. (**a**) Schematic representation of tumor rechallenge assay, in which human primary CD8^+^ and CD4^+^ CAR expressing T cells mixed in a 1:1 ratio were stimulated every 2 days with NALM6 cells at an E:T ratio of 1:10. (**b**) T cells and (**c**) NALM6 cells were quantified by FACS at each time point prior to restimulation, along with (**d**) T cell memory differentiation markers (n = 3 technical replicates representative of 2 biological replicates). *P* values in panel (**b**) are 0.004396, 0.001890, 0.010757, 0.000033, 0.000060, 0.000003, 0.000652, 0.003049, and 0.000080 for Var1 vs. BB**ζ** on days 5, 7, 10, 12, 14, 17, 19, 21, and 24 of rechallenge for CD8+ CAR-T cells. *P* values are 0.000340, 0.010693, 0.000019, 0.000537, 0.000374, and 0.000409 for days 3, 7, 17, 19, 21, and 24 of rechallenge for CD4+ CAR-T cell expansion. *P* values for NALM6 cell expansion in panel (**c**) are 0.002063, 0.005311, 0.000043, 0.000790, and 0.000589 for Var1 vs. BB**ζ** on days 7, 10, 12, 21, and 24 of rechallenge. *P* values for panel (**d**) are 0.011291, 0.000865, 0.000019, and 0.000014 for Var1 vs. BB**ζ** for days 4, 6, 11, and 18 of rechallenge. All *P* values for panels (**b**)-(**d**) were determined by multiple two-sided unpaired student’s t tests with Benjamini-Hochberg correction (df = 4 for all). Data shown are means ± s.e.m. (**e**) Exhaustion marker expression of CD4^+^ and CD8^+^ CAR-T cells on day 22 of rechallenge (n = 3 technical replicates representative of 2 biological replicates). *P* values are 0.0001 for CD4^+^ Var1 vs. BB**ζ**, 0.0041 for CD4^+^ Var3 vs. BB**ζ**, 0.0028 for CD4^+^ Var1 vs. Var3, 0.0004 for CD8^+^ Var1 vs. BB**ζ**, 0.0080 for Var3 vs. BB**ζ**, and 0.0004 for CD8^+^ Var1 vs. Var3 for PD-1. *P* values are 0.0019 for CD4^+^ Var1 vs. BB**ζ**, 0.0042 for CD4^+^ Var3 vs. BB**ζ**, 0.0017 for CD8^+^ Var1 vs. BB**ζ**, 0.0034 for CD8^+^ Var3 vs. BB**ζ**, and 0.0386 for CD8^+^ Var1 vs. Var3 for TIM3. *P* values were determined using a two-tailed unpaired student’s t-test (df = 4). Data shown are individual points along with means ± s.e.m.

### Var1 induces a unique transcriptional profile associated with T cell activation and persistence

Given that the variant CARs exhibited unique functional phenotypes relative to BB**ζ** *in vitro*, we asked whether the signaling perturbations introduced by the ICD combinations in Var1 and Var3 produced unique transcriptional programs in response to tumor challenge. We subjected primary CD4^+^ and CD8^+^ T cells expressing each construct to three high-burden NALM6 challenges at an E:T ratio of 1:10 over a five-day period (on days 0, 3, and 5) before performing single-cell RNA-sequencing (scRNA-seq) 48 hours after the third tumor challenge. Dimensionality reduction and unsupervised clustering revealed five transcriptionally distinct cell clusters (**Fig. 5a**). While CD4^+^ and CD8^+^ cells cluster together to some extent (**Supplementary Fig. 9**), the separation between clusters was mainly defined by CAR variant, with Var1-expressing cells confined primarily to C0 and C1, and BB**ζ**- and Var3-expressing cells localized to C2, C3, and C4 (**Fig. 5a**); notably, all CAR-expressing cells also clustered separately from unstimulated, untransduced controls (**Supplementary Fig. 10**). Chi-squared analysis confirmed significant enrichment of Var1 cells in C0 and C1, while the distribution of BB**ζ** and Var3 cells were indistinguishable (**Fig 5b**), suggesting that the Var1 construct drives a unique transcriptional phenotype following repeated high-burden tumor challenge.

**Figure 5.**
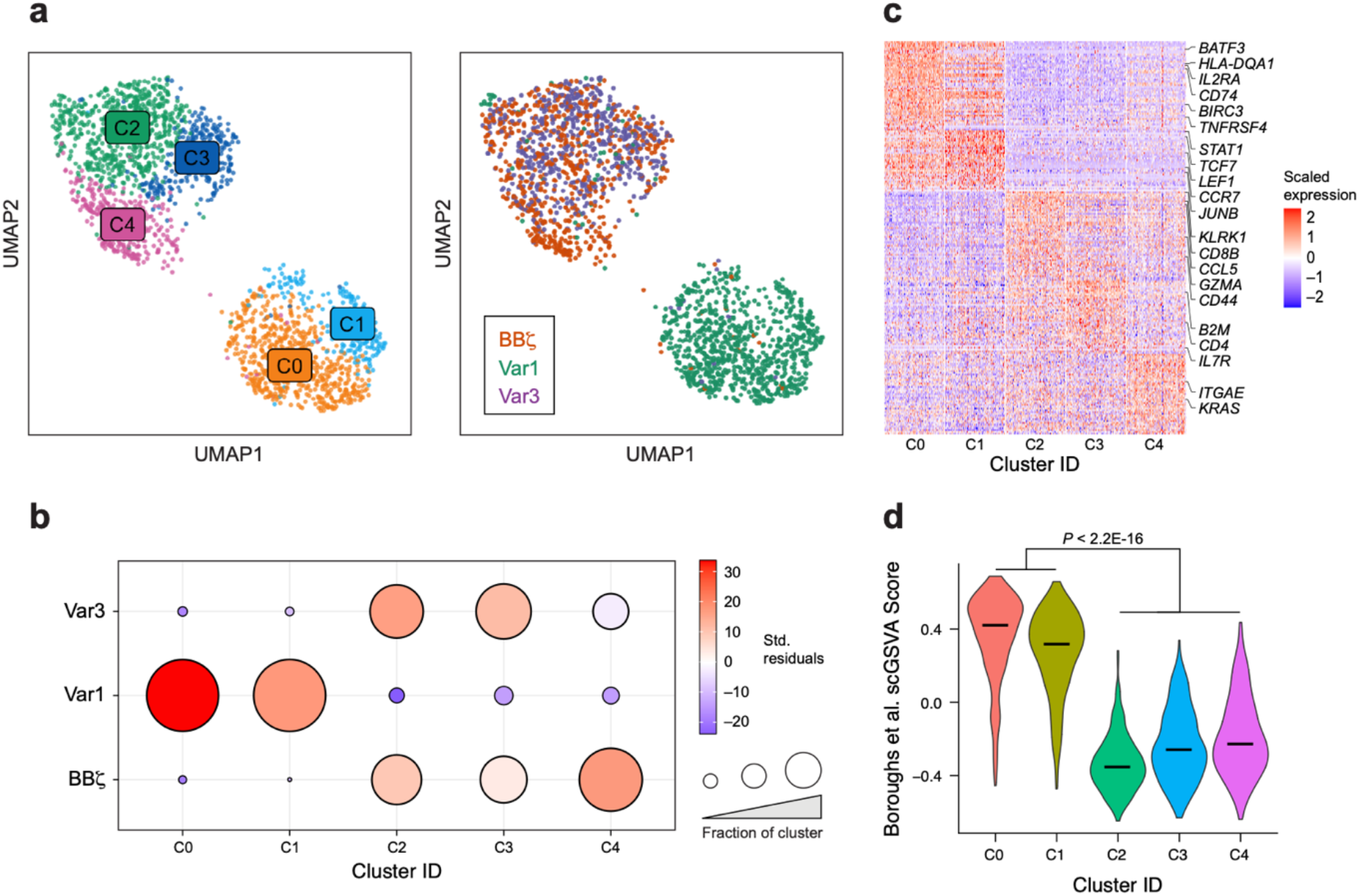
Var1 upregulates transcription of genes involved in T cell persistence following tumor rechallenge. (**a**) UMAP embeddings of merged scRNA-seq profiles following rechallenge of human primary CD8^+^ and CD4^+^ CAR T cells (mixed in a 1:1 ratio) with NALM6 cells three times at an E:T ratio of 1:10 colored by cell state (left) and CAR identity (right) (n = 2,082 cells). (**b**) Chi-square enrichment values for each CAR candidate within each cluster, represented by the Pearson residuals measuring the difference between the observed and expected CAR frequencies within each cluster (df = 8). (**c**) Heat map representing the normalized expression of the top 50 differentially expressed genes within each cluster, as determined by Wilcoxon-Rank Sum test with Bonferroni correction. (**d**) Single-cell gene set variation analysis (scGSVA) scores measuring enrichment of a previously published BB**ζ**-specific gene set ^33,46^ within each T cell cluster. Bars represent median scGSVA values. *P* values were determined using a Wilcoxon-Rank Sum test.

To characterize the transcriptional program associated with Var1 signaling, we performed differential expression analysis to define cluster-specific markers (**Fig 5c** and **Supplementary Table 2**). Among the most significantly overexpressed genes in both C0 and C1 were several genes related to T cell activation (*IL2RA, TNFRSF4*), including a broad repertoire of MHC class II-related genes (**Supplementary Fig. 11a**). In addition, we observed significant upregulation of memory markers such as *CCR7*, as well as genes within pathways thought to promote T cell persistence and memory, including those related to non-canonical NF-kB signaling (*BIRC3*, *TRAF1, NFKB2*)^28,29^ and AP-1 transcription factors (*BATF3, JUNB*)^30^ (**Supplementary Fig. 11b**). Although the transcriptional profiles defined by C0 and C1 were largely similar, cells within C1 uniquely overexpressed *TCF7* and *LEF1*, which encode for transcription factors thought to be important for T cell stemness and memory (**Supplementary Fig. 11c**).^31^ Similar hits were observed when directly comparing average expression profiles of Var1 cells to BB**ζ** (**Supplementary Fig. 12 and Supplementary Table 3**). Considering these differences at the gene level, we sought to characterize Var1-associated programs at the pathway level; we employed single-cell gene set variation analysis (scGSVA) to assign scores to each cell within the dataset on the basis of their relative expression of genes within canonical and curated T cell gene sets (**Supplementary Table 4**).^32^ Interestingly, one of the most significantly upregulated pathways among Var1-expressing cells was a geneset recently reported to distinguish 19BB**ζ** CARs from 1928**ζ** (**Fig. 5d**),^33^ suggesting that Var1 might induce similar transcriptional programs to that of BB**ζ** but to a greater extent. Indeed, orthogonal pathways with minimal gene overlap that have been separately demonstrated to distinguish BB**ζ** CARs are similarly enriched in C0 and C1 (**Supplementary Fig. 13**).^28,33^ Taken together, these results suggest that Var1 triggers a coordinated transcriptional response that promotes enhanced persistence and long-lived memory formation relative to BB**ζ** in response to antigen stimulation.

### Var1 shows comparable activity to that of a BBζ CAR in a leukemia mouse model

We next tested whether our novel CARs produced distinct *in vivo* outcomes in the context of a xenograft mouse model of B cell leukemia. We injected NOD/SCID/IL2R^null^ (NSG) mice with 5×10^5^ luciferase-expressing (FLuc^+^) NALM6 cells intravenously followed by treatment with a 1:1 mixture of untransduced or CAR-expressing CD4^+^ and CD8^+^ T cells 4 days later (**Fig. 6a**). When treated with 2×10^5^ CARs, Var1-treated mice demonstrated slightly delayed tumor outgrowth and a subtle but not statistically significant increase in survival compared to BB**ζ**-treated mice, with no discernible signs of toxicity (**Fig. 6b-d and Supplementary Fig. 14-15**). Var3-treated mice showed comparable tumor progression relative to BB**ζ**-treated mice. Upon collecting peripheral blood on day 27, we detected significantly elevated levels of EGFP^+^ CAR T cells in Var3-treated mice relative to BB**ζ**-treated mice, while the number of circulating Var1 CARs was significantly lower despite producing a similar survival benefit (**Fig. 6e**). All groups of mice exhibited comparable levels of CD19^+^ NALM6 cells (**Fig. 6f**). We additionally treated mice with a higher dose of 1×10^6^ CARs. While we observed sustained remission across all groups, we noted that Var1-treated mice showed delayed tumor control in this paradigm (**Supplementary Fig. 16-17**).

**Figure 6.**
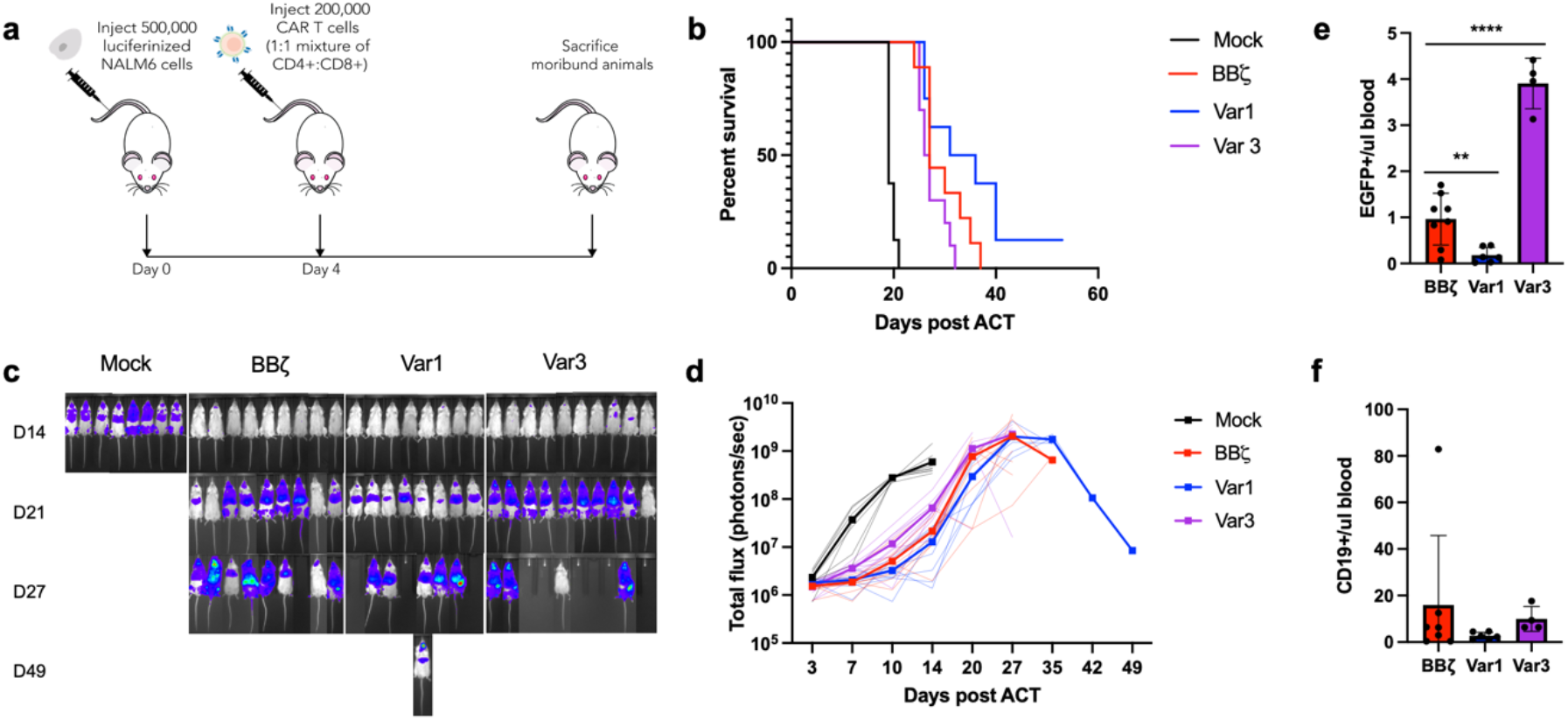
Variant 1 demonstrates similar tumor control to that of a BBζ CAR *in vivo*. (**a**) Experimental design: NSG mice were intravenously infused with 5×10^5^ FLuc^+^ CD19^+^ NALM6 cells, then treated with 2×10^5^ mixed CD4^+^ and CD8^+^ (1:1) CAR T cells or untransduced control T cells (n = 8 mice for untransduced, 9 for BB**ζ**, 8 for Var1, and 10 for Var3). (**a**) Kaplan-Meier curve for overall survival. *P* values, as determined using a Log-rank test, are 0.1268 for Var1 vs. BB**ζ** and 0.0788 for Var3 vs. BB**ζ** (df = 1). Tumor bioluminescence was assessed every 3-7 days by imaging for luciferase activity. (**c**) Representative images and (**d**) quantification of total photon counts are shown. Data shown are individual points along with means in bold. (**e**) Concentrations of EGFP^+^ CAR T cells and (**d**) CD19^+^ NALM6 cells in the peripheral blood were determined on day 27 after ACT by FACS (n = 8 mice for BB**ζ**, 6 for Var1, and 4 for Var3). *P* values panel (**e**) are 0.0067 for Var1 vs. BB**ζ** (df = 12) and <0.0001 for Var3 vs. BB**ζ** (df = 10). *P* values were determined using a two-tailed unpaired student’s t test. Data shown are individual points along with means ± s.e.m.

## Discussion

Despite the clinical promise of CAR-T therapies, efforts to improve CAR-T function by systematically optimizing CAR signaling remain limited. A previous effort successfully identified functionally signaling CARs from a pool of signaling domains, but was significantly smaller in scope, had a relatively high false positive rate, and lacked the sequencing analyses necessary to comprehensively examine any selected variants.^34^ Here, we describe CARPOOL, a library-based functional screening platform for rapidly enriching and identifying novel CARs with clinically useful phenotypes. As a proof of principle, we constructed a 700,000-member CD19 CAR library with diverse ICD combinations to discover novel CARs based on their ability to robustly activate Jurkat T cells. We believe our observations from our selections (**Fig. 1d**) are representative of many library-based screening approaches: essentially every CAR included at least one ITAM domain and many contained costimulatory domains, which matches the known rules of CAR design. However, the selection, covariation, and spatial orientations of each selected construct could not have been predicted from first principles.

When we validated selected CARs, we found that Var1 CARs showed enhanced anti-tumor activities *in vitro* and comparable tumor control *in vivo* compared to those of the BB**ζ** CAR currently being used in the clinic. It should be noted that the CD40 and CD3ε ICDs that comprise the two membrane-proximal ICDs of Var1 were separately reported to enhance CAR-T cell function, with CD40 augmenting CAR-T effector function and MyD88 and CD3ε promoting CAR-T cell persistence through Csk and p85 recruitment via the ITAM and basic residue rich sequence (BRS), respectively.^5,7,35^ Additionally, DAP12, the membrane distal ICD of Var1, has been demonstrated as a functional ICD in both CAR-T and CAR-NK cells.^18,19^ While each of these domains has been examined in isolation, this is the first time they have been collectively characterized. Further work may uncover the extent to which these signaling inputs synergize with each other to generate the observed T cell phenotypes.

Two key considerations for contextualizing our results are the translation between Jurkat and primary T cells, and translation to *in vivo* settings. While our study demonstrated that selections conducted in Jurkat cells can reliably identify novel CARs, there are inherent limitations to the effector functions that can be selected for due to the physiology of the cell line. Being an immortalized cell line derived from T cell leukemia, Jurkats continually divide, exhibit altered basal signaling and metabolism, and are not cytotoxic. ^36,37^ Therefore, there are desirable T cell functions such as resistance to exhaustion and efficient tumor killing that cannot be directly selected for in a Jurkat-based library. The fact that Var1 exhibited many of these phenotypes despite being selected in Jurkats indicates either these functions correlate with features that can be selected for in Jurkat cells, or that there are a wide range of emergent properties accessible simply by altering signaling inputs.

Additionally, we consider the differences in results that were observed *in vitro* compared to those observed *in vivo*. Var1 showed superior function in many *in vitro* assays while appearing comparable but not significantly superior *in vivo*. Discrepancies in translation between the two have been well reported.^38–40^ These differences are notable in that several of the differences observed *in vitro* involved either the production of cytokines that would not have function in a xenograft mouse model, or persistence improvements that may only be relevant upon consistent antigen exposure. It is therefore possible that the comparable results between Var1 and 19BB**ζ** are due to commonalities in CAR function or limitations of the *in vivo* model itself.

Extending the CARPOOL system for selection in primary T cells would make a broader range of phenotypes available for use as selection criteria. Doing so could enable the rapid identification of novel CAR clones that could address existing challenges in translating CAR-T cell therapies to solid tumor indications, such as lack of persistence due to tumor-mediated immunosuppression and inefficient trafficking and tumor infiltration ^2,41–43^. This may require more specialized selection strategies, such as employing T cell surface markers indicative of cytotoxicity, long-lived memory differentiation, and tumor infiltration, in addition to introducing selection conditions that mimic the microenvironment of solid tumors. CARPOOL could also be adapted to systemically optimize CAR ICD designs for other emerging immune cell therapy modalities such as CAR-NK cells and CAR-macrophages ^44,45^. In summary, CARPOOL presents a versatile, streamlined method for functionally engineering synthetic receptors for use in immune cell therapies.

## Methods

### Cell lines

HEK293T (CRL-3216) and Clone E6-1 Jurkat (TIB-152) lines were purchased from ATCC, while Clone G5 NALM6 (CRL-3273) cells were purchased from ATCC and transduced to stably express firefly luciferase (FLuc) along with a puromycin resistance cassette. Cell lines were routinely mycoplasma-tested using the MycoAlert PLUS Mycoplasma Detection Kit (Lonza).

### Plasmid construction

The plasmid pHIV-EGFP was gifted by Bryan Welm & Zena Werb (Addgene plasmid #21373) and pMD2.G and psPAX2 were gifted by Didier Trono (Addgene plasmid #12259 and #12260). To generate 2nd generation CD19 CAR-EGFP plasmid, a codon optimized gene encoding CD19 CAR composed of Myc-epitope tagged FMC63 scFv, IgG4 hinge, CD28 transmembrane domain, and intracellular domains derived from human 4-1BB and CD3**ζ** was PCR amplified from geneblocks purchased from IDT and cloned into the 3nd generation lentiviral vector pHIV-EGFP using Gibson Assembly. In order to generate a backbone vector for CAR plasmid library, the Myc epitope tag from CD19 CAR-EGFP plasmid was changed to a Flag epitope tag, and six tyrosine residues from CD3**ζ** ITAM domains were mutated into phenylalanines to prevent any unmodified yet functional CARs in the library from contaminating library selections. The signaling diversified CAR plasmid library was generated by PCR amplification of each intracellular domain (**Supplementary Table 1**) at each of the 3 positions, with the forward and reverse primers adding unique linkers for each position. These products were then pooled at equimolar ratios for each position and combined with a pool of randomized 18mer barcode sequences for overlap extension PCR. These were then inserted into degenerate CD19 CAR backbone vector at PacI and BamHI restriction enzyme sites to replace the tyrosine mutated BB**ζ** intracellular signaling components via Gibson Assembly. Final products were electroporated into DH10B electrocompetent E.coli cells (Thermo Scientific, EC0113) and purified to achieve a highly diverse plasmid library.

### Lentiviral production

Lentiviruses were generated by first transfecting 70% confluent HEKs with transfer plasmid, pMD2.g (VSVg), and psPAX2 combined at a plasmid mass ratio of 24:1:3 that was complexed with PEI at a DNA:PEI mass ratio of 1:3. For a confluent T225 flask, 60 ug of transfer plasmid was used for transfection. Media was changed 3-6 hours after transfection and lentiviral particles were harvested in the supernatant 48-120 hours after transfection. The supernatant was then filtered through a 0.45 um low protein binding filter, and centrifuged for 1.5 hours at 100,000x g. The pellet was then resuspended in serum-free OptiMEM overnight at 4°C and stored at −80°C.

### Human T cell activation, transduction, and expansion

Peripheral blood mononuclear cells from healthy donors were purified from buffy coats purchased from Research Blood Components or leukopaks purchased from Stem Cell Technologies using Ficoll-Paque PLUS (GE Healthcare) density gradient centrifugation with SepMate tubes (Stem Cell Technologies) as per manufacturer instructions. Primary CD4+ or CD8+ T cells were isolated using EasySep Human CD4+ or CD8+ T Cell Enrichment Kits (Stem Cell Technologies) and cultured in RPMI 1640 (ATCC) supplemented with 10% fetal bovine serum, 100 U/ml penicillin-streptomycin (Corning), 100 IU/ml recombinant human IL-2 (R&D Systems), and 50 μM beta-mercaptoethanol (Fisher). Prior to transduction, T cells were activated using a 1:1 ratio of DynaBeads Human T-Activator CD3/CD28 (Thermo Fisher) for 24 hours, after which 8 μg/mL of polybrene (Santa Cruz Biotechnology) and concentrated lentivirus were added to culture at a multiplicity of infection of 10 for single lentiviral constructs and 0.5 for pooled library encoding lentivirus. After 3 days, DynaBeads and lentivirus were removed and cells were sorted for EGFP using a BD FACSAria II. Cells were rested for 4 days prior to characterization and maintained at a density of 5×10^5^-2×10^6^ cells/ml throughout this process.

### Flow cytometry and cell sorting

Cells were washed with 1X PBS (Sigma) supplemented with 0.5% bovine serum albumin (RPI) and 2 mM EDTA, then surface stained by incubating with antibodies for 15 minutes on ice. They were subsequently washed again prior to flow analysis on a BD Accuri C6 or Beckman Cytoflex S or cell sorting with a BD FACSAria II or Sony MA900. Anti-CD4 (clone SK3), anti-CD8 (clone SK1), anti-PD-1 (clone EH12.2H7), anti-TIM3 (clone F38-2E2), anti-LAG3 (clone 11C3C65), anti-CD3 (clone OKT3), anti-CD62L (clone DREG-56), anti-CD45RA (clone HI100), and anti-CD69 (clone FN50) antibodies were purchased from Biolegend. Anti-Myc (clone 9B11) and anti-Flag (D6W5B) antibodies were from Cell Signaling Technology.

### CAR-T functional selections

In preparation for selections, 1×10^8^ Jurkat T cells were transduced with lentivirus at an MOI of 0.5 with 8 ug/ml polybrene (Santa Cruz Biotechnology) and spinfected at 1000x g for 1.5 hours at 32°C. Virus was removed after 2 days of transduction and the cells were sorted the following day for EGFP, with 20x library coverage being maintained based upon the theoretical maximm diversity from the previous round previously throughout this process. For a round of selection, cells were stimulated with 1 or 10 nM rhCD19 for 4-6 hours as indicated, then stained for CD69 or CD69 and PD-1 expression. The top 5% T cells as measured by CD69^high^ or CD69^high^PD-1^low^ were sorted on a BD FACS Aria II (with at least 5×10^5^ cells being collected). Cells were then rested without antigen and expanded for 7 days before subsequent rounds of selection. After each round of selection, at least 1×10^6^ cells were sampled for NGS sequencing (whereas 6×10^7^ and 2×10^7^ cells were sampled for unselected and EGFP sorted groups, respectively). NGS sequencing data was deconvoluted and analyzed using a custom package called DomainSeq, as described in the manuscript.

### In vitro rechallenge assay

CAR-transduced CD4^+^ and CD8^+^ T cells (50,000 cells each) were mixed at a 1:1 ratio, then co-cultured with target NALM6 cells at an effector to target (E:T) ratio of 1:10 in IL-2 deficient media. Every 2-3 days, approximately 10% of the culture volume was taken out for flow analysis and stained with antibodies targeting CD4, CD8, CD62L, and CD45RA. Then, 100,000 CAR-T cells were taken out from the original culture and re-plated with a fresh batch of NALM6 cells at a 1:10 E:T ratio. CAR T cells were sampled for scRNA-seq analysis at day 7, which was 48 hours following the third NALM6 challenge. On day 22, cells were also stained for exhaustion markers (PD-1, TIM3, and LAG3).

### Cytotoxicity assay

NALM6 cells expressing firefly luciferase (FLuc) were co-cultured with CD8^+^ T cells for 24 hours in IL-2 deficient media at various E:T ratios. Cells were then harvested and washed prior to cell lysis and addition of luciferin substrate from the Bright-Glo Luciferase Assay System (Promega). The resulting luminescent signal was measured using a Tecan Infinite M200 Pro. Signals were normalized to negative controls containing only target cells.

### Cytokine secretion assay

Following stimulation of human primary CAR-T cells with NALM6 cells, the concentrations of human IL-2 and IFN-γ were measured using a IL-2 Human Uncoated ELISA Kit (Thermo Fisher) and IFN-γ Human Uncoated ELISA Kit (Invitrogen), respectively. The resulting signal was measured on a Tecan Infinite M200 Pro plate reader, and the concentrations were determined by comparison to known standards per the manufacturer’s instructions. Polyfunctional cytokine and chemokine secretion profiles in response to tumor challenge were determined using the 41-plex MILLIPLEX MAP Human Cytokine/Chemokine Magnetic Bead Panel from Miltenyi and measured on a Luminex FlexMap 3D system.

### PacBio and Illumina sequencing

Genomic DNA from selected cells was purified using the PureLink Genomic DNA Mini Kit (Thermo Fisher). For PacBio sequencing, PCR amplicons encoding CAR signaling domains and barcode regions were attached with SMRTbell adaptors using the SMRTbell Template Prep Kit 1.0 (Pacific Biosciences) and sequenced using a PacBio Sequel system. For Illumina sequencing, barcode regions were PCR amplified to conjugate P5 and P7 adaptor sequences and sequenced on an Illumina MISeq system.

### Single-cell sequencing

19BB**ζ**, Var1, and Var3-expressing CAR T cells were sampled from NALM6 cocultures, as well as untransduced, unstimulated T cells. Separately, samples were enriched for live cells using a Dead Cell Removal (Annexin V) Kit (Stem Cell Technologies), then labeled with unique anti-human TotalSeq-B hashing antibodies (BioLegend). Following labeling, approximately 2,500 cells from each of the four samples were pooled before encapsulation in a single channel of the Chromium Single Cell 3’ v3.1 platform (10X Genomics). Gene expression (GEX) libraries were constructed based on manufacturer instructions, while hashing antibody libraries were constructed as reported previously.^46^ The resulting libraries were pooled at a 1:10 ratio of antibody-to-GEX before sequencing on an Illumina NextSeq500 to a depth of 60,000 reads per cell. Reads were aligned to the Genome Reference Consortium Human Build 38 (GRCh38), and a cell-gene matrix was generated using the CellRanger pipeline (10X Genomics; v4.0.0). Downstream analysis was performed using the Seurat package (v4.0.0).^46,47^ In brief, cells were first assigned sample identity based on the detection of a single hashing antibody following normalization of antibody reads using the HTODemux algorithm.^46^ Next, low-quality cells were filtered out on the basis of mitochondrial reads (>25%). Filtered data for each cell was normalized to total expression, and cell cycle-related genes were regressed out using the ScaleData function. To identify distinct transcriptional states, linear dimensionality reduction was performed on the scaled, normalized data, followed by shared nearest neighbors clustering on the basis of the first 40 principal components. Differentially expressed genes, both within clusters and across samples, were identified by a Wilcoxon-Rank Sum test between the populations of interest. For pathway-level analyses, individual cells were assigned scGSVA scores on the basis of their relative expression of genes within all canonical pathways, immunologic signature gene sets (Broad Institute; C2 and C7 gene sets, respectively), or T cell-specific curated pathways (**Supplementary Table 4**), as previously described. ^32^

### Xenogeneic mouse models

All animal studies were performed in accordance with guidelines approved by the MIT Division of Comparative Medicine and MIT Committee on Animal Care (Institutional Animal Care and Use Committee). Male NOD/SCID/IL2R^null^ (NSG) mice were purchased from Jackson Laboratory and housed in the animal facilities at MIT. At age 8-12 weeks old, mice were injected intravenously via the tail vein with 5×10^5^ FLuc^+^ NALM6 cells. CD4+ and CD8+ T cells were prepared separately as described above, then sorted for EGFP on the day of DynaBead removal; after 4 days of rest, they were then mixed at a 1:1 ratio. Mice were then treated with 2×10^5^ or 1×10^6^ CAR-T cells or untransduced control T cells intravenously via the tail vein 4 days after NALM6 injection. Tumor progression was subsequently monitored every 3-7 days using the IVIS Spectrum imaging system (PerkinElmer) to measure bioluminescent signal after intraperitoneal administration of 0.15 mg of luciferin substrate per gram of body weight (PerkinElmer 122799). Total photon counts were quantified using LivingImage software. Mice were monitored daily and euthanized upon observing signs of discomfort or morbidity, graft versus host disease, or as recommended by the veterinarian. Where indicated, peripheral blood was collected to measure T cell expansion and persistence by flow cytometry. Red blood cells were lysed from the collected tissues using ACK Lysing Buffer (Thermo Fisher A1049201) and washed with 1X PBS supplemented with 0.5% BSA and 2 mM EDTA prior to antibody staining and FACS analysis.

### Statistical analysis

Statistical analyses were performed using the Prism 9 (GraphPad) software, with the exception of the single-cell sequencing data which was analyzed in R Studio using base packages or those described above. Sample sizes were not predetermined using statistical methods. For statistical comparisons between two groups, significance was determined using two-tailed unpaired parametric t-tests or nonparametric Wilcoxon Rank Sum tests. For *in vivo* experiments, differences in overall survival were analyzed using a log-rank test and displayed in a Kaplan-Meier curve. Adjusted *P* values < 0.05 after multiple hypothesis correction, where required, were considered statistically significant. The statistical test used for each experiment is noted in the relevant figure legend.

### Reporting Summary

Further information regarding study design is available in the Nature Research Reporting Summary appended to this article.

### Data Availability

The NGS datasets have been deposited in the Sequence Read Archive and are available under accession number PRJNA744269, while the scRNA-seq data has been deposited in the Gene Expression Omnibus under accession number GSE179767. The DomainSeq-processed CARPOOL selection data is available in the GitHub repository at https://github.com/birnbaumlab/Kyung-et-al-2021. All data generated or analyzed during this study are included in this published article and its supplementary information files.

### Code Availability

The code used to analyze the domain composition of selected CARs can be accessed in the DomainSeq repository at https://github.com/birnbaumlab/Kyung-et-al-2021.

## Acknowledgements

We would like to thank the Koch Institute Swanson Biotechnology Center for their technical support, especially the Flow Cytometry Facility, Animal Facility, Animal Imaging and Preclinical Testing Facility, MIT BioMicro Center, and High Throughput Sciences Facility. We also thank G.A. Paradis, P. Chamberlain, H. Holcombe, V. Spanoudaki, S.S. Levine, and B.A. Joughin for many helpful discussions and suggestions. We also thank Y.Y Chen for providing the sequence for the 19BB**ζ** CAR.

This work was funded by grants from the Packard Foundation, the Pew Foundation, the Deshpande Center, and the National Institutes of Health to M.E.B., and a Fellowship from the Human Frontier Science Program to T.K. This work was also supported in part by the Mark Foundation, the NIH (award CA247632), and the Bridge Project, a partnership between the Koch Institute for Integrative Cancer Research at MIT and the Dana-Farber/Harvard Cancer Center, along with award Number T32GM007753 from the National Institute of General Medical Sciences. The content is solely the responsibility of the authors and does not necessarily represent the official views of the National Institute of General Medical Sciences or the National Institutes of Health. This research is supported in part by the National Research Foundation, Prime Minister’s Office, Singapore under its Campus for Research Excellence and Technological Enterprise (CREATE) programme, through Singapore MIT Alliance for Research and Technology (SMART): Critical Analytics for Manufacturing Personalised-Medicine (CAMP) Inter-Disciplinary Research Group.

## Author information

### Contributions

Conception of project: T.K, M.E.B. Conducting experiments: T.K., K.S.G., C.R.P, A.R., A.Q.Z., Y.L., C.K., A.S. Data analysis: T.K., K.S.G., C.R.P, P.V.H, A.R., B.J. Supervision: D.A.L., M.T.H., D.J.I., M.E.B. Writing manuscript: T.K., K.S.G., C.R.P., M.E.B. Editing manuscript: All authors

## Ethics declarations

The library approach described in this manuscript is the subject of a US patent application (US20200325241A1) with T.K. and M.E.B. as inventors. M.E.B. is a founder, consultant, and equity holder of Viralogic Therapeutics and Abata Therapeutics. T.K. is presently an employee of Catamaran Bio.

