## Supplementary Figures 1-17 for "CARPOOL: A library-based platform to rapidly identify next generation chimeric antigen receptors"

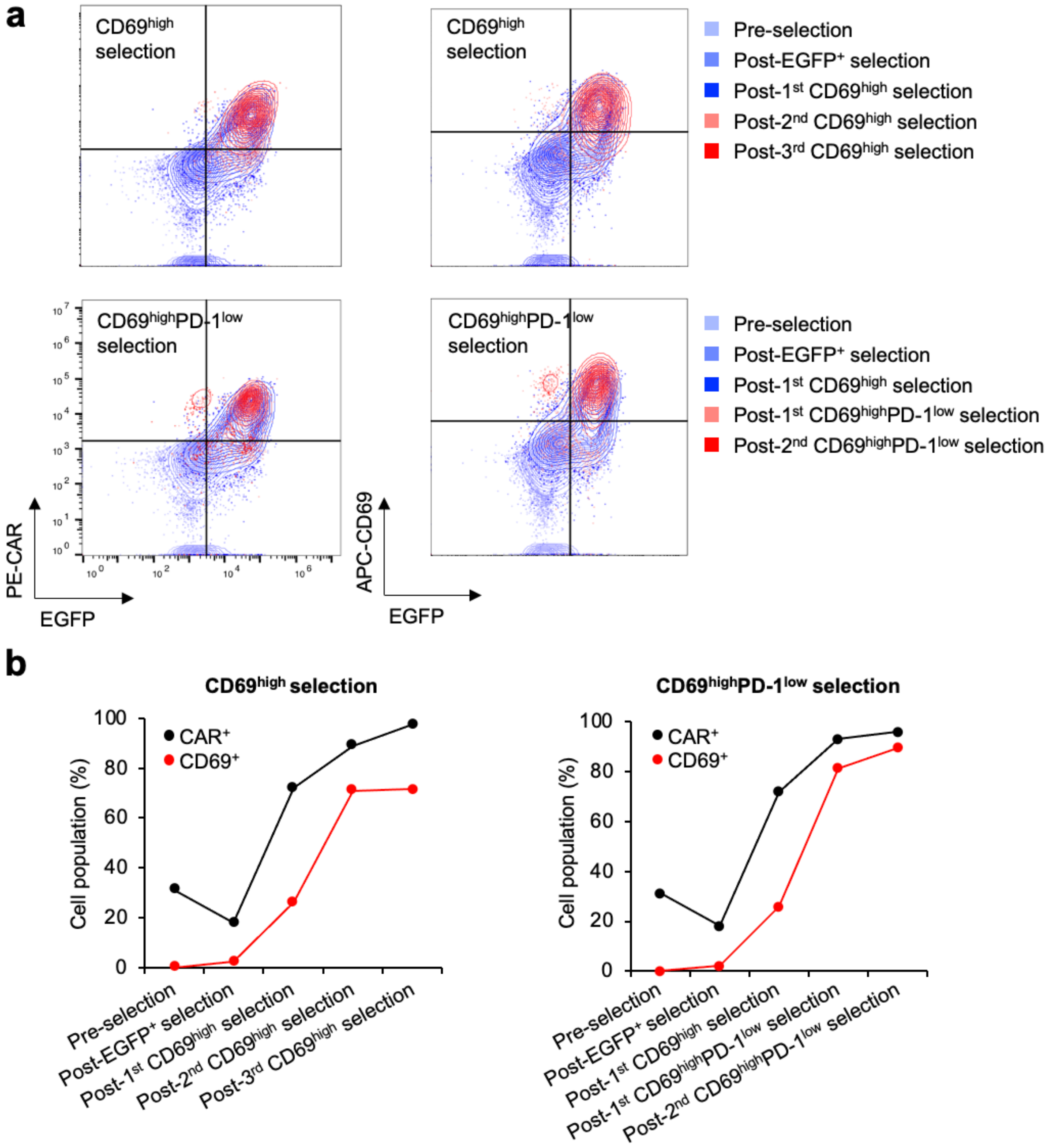


**Supplementary Figure 1. Sorting for CD69 expression enriches CD69$^{high}$ populations**
(**a**) Flow cytometry plots showing increase in transgene (EGFP$^{+}$) and CD69 expression upon rounds of CAR stimulation and selection. (**b**) Quantitation of CAR and CD69 expression.

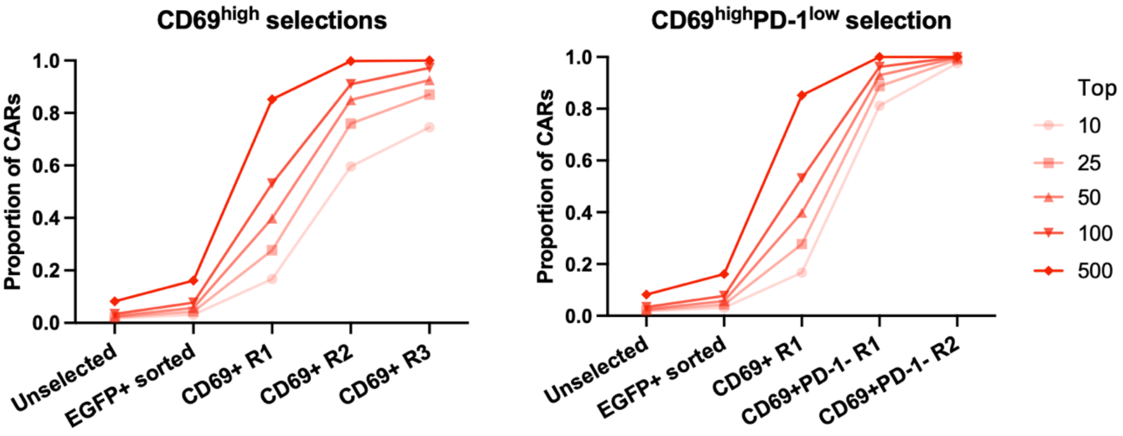


**Supplementary Figure 2.** Proportions of most enriched fractions of barcodes from each round of selection.

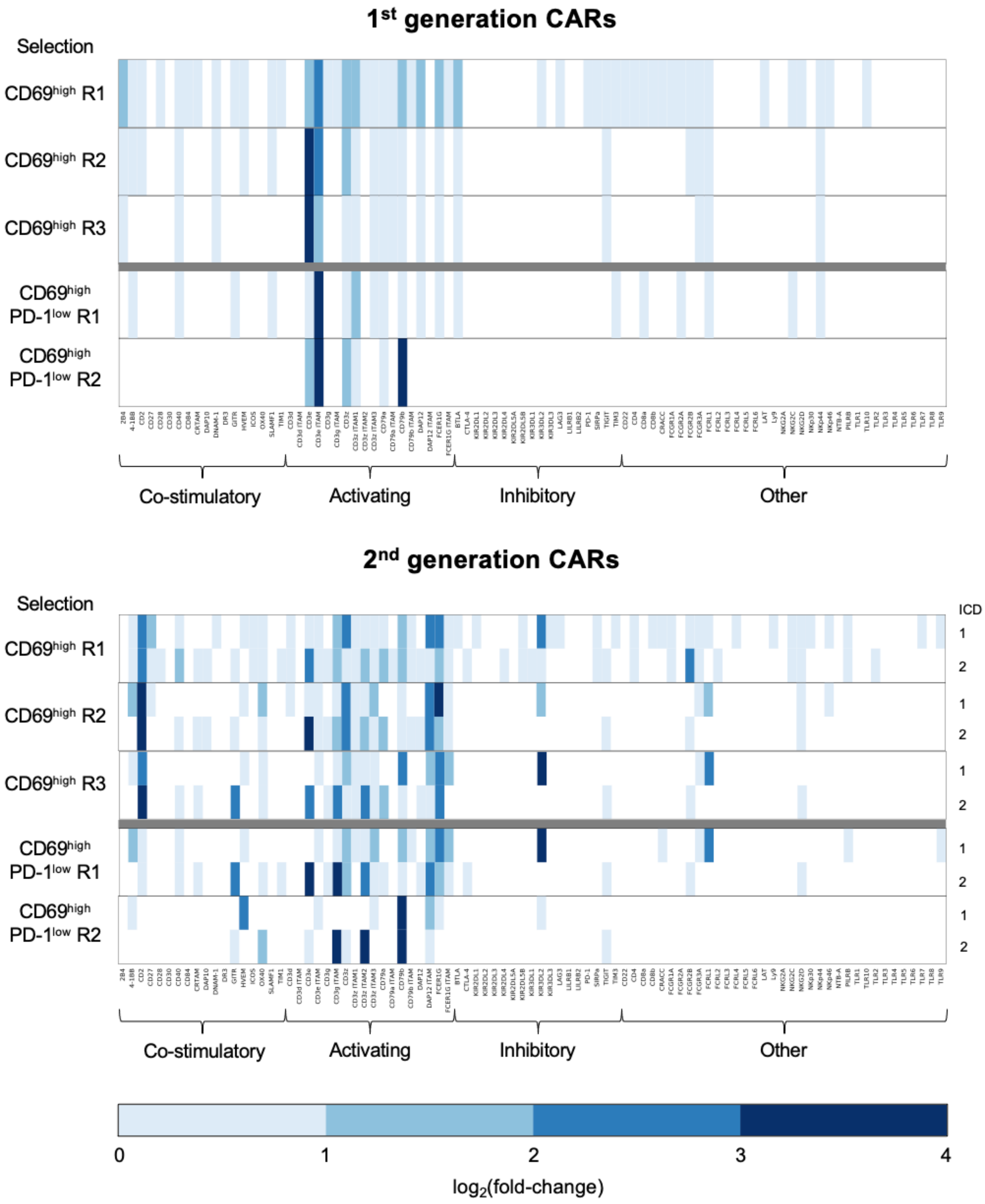


**Supplementary Figure 3.** Heatmap representing $log_2$(fold-change) of individual ICDs found in (**a**) 1st and (**b**) 2nd generation CARs throughout rounds of selection, oriented by ICD position within the CAR intracellular domain relative to the plasma membrane.

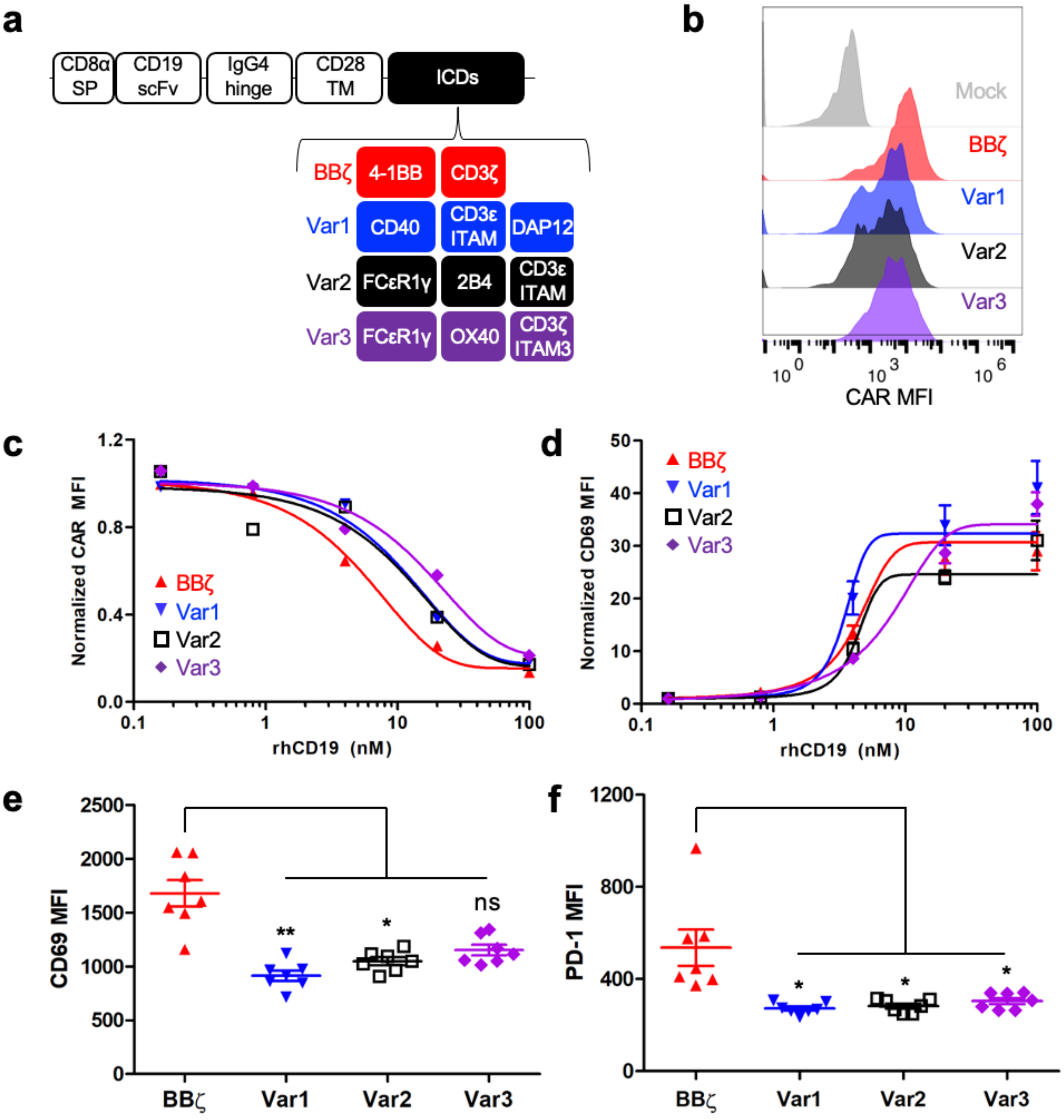


**Supplementary Figure 4.** Characterization of fundamental aspects of CAR variant expression in Jurkat T cells. (**a**) Design of CD19 targeted CAR candidates. (**b**) CAR surface expression (n = 1), (**c**) internalization (n = 1), and (**d**) CD69 upregulation upon antigen stimulation with recombinant human CD19 (n = 6 technical replicates representative of 2 biological replicates). (**e**) Basal CD69 (activation) and (**f**) PD-1 expression of unstimulated CAR-T cells (n = 7 technical replicates representative of 2 biological replicates). *P* values shown in panel (**e**) are 0.002 for Var1 vs. BB**ζ** and 0.0149 for Var2 vs. BB**ζ**. *P* values shown in panel (**f**) are 0.0117 for Var1 vs. BB**ζ**, 0.0110 for Var2 vs. BB**ζ**, and 0.0192 for Var3 vs. BB**ζ**, as determined by two-tailed unpaired student's t tests (df = 12). Data shown are individual points in panel (**c**) as well as means ± s.e.m in (**d**)-(**f**).

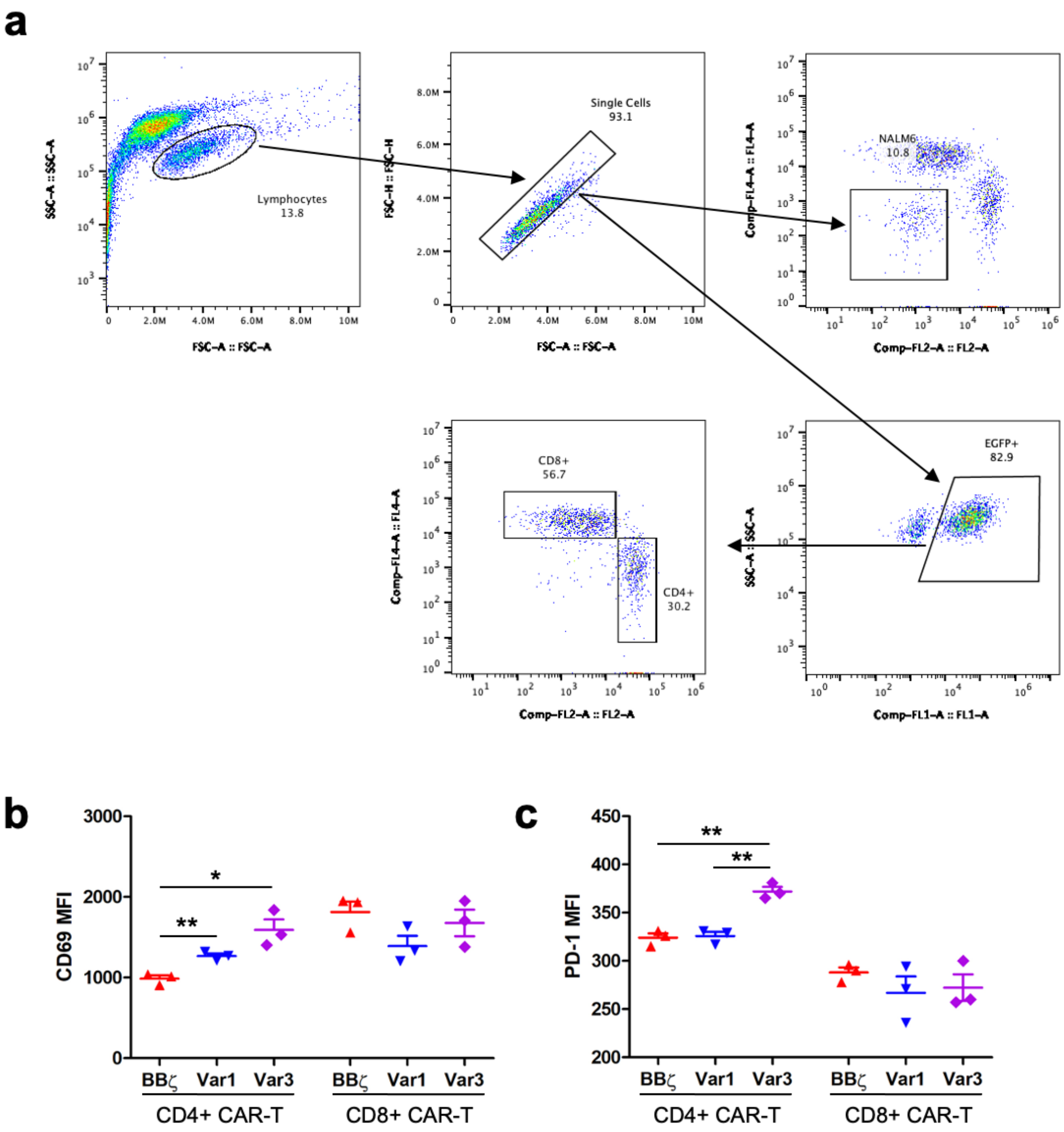


**Supplementary Figure 5.** Representations demonstrating basal phenotypic status of CAR-T cells in human primary CD4$^+$ and CD8$^+$ T cells. (**a**) FACS plots showing gating strategy. (**b**) Basal CD69 and (**c**) PD-1 expression in unstimulated human primary CD4$^+$ and CD8$^+$ CAR-T cells (n = 3 technical replicates representative of 2 biological replicates). *P* values for CD69 expression are 0.0050 for Var1 vs. BB**ζ** and 0.0108 for Var3 vs. BB**ζ** in CD8$^+$ T cells. *P* values for PD-1 expression are 0.0020 for Var3 vs. BB**ζ** and 0.0020 for Var1 vs. Var3. (**d**) FACS plots showing CAR expression vs. CD69 expression in unstimulated CAR-T. (**e**) FACS plots showing memory phenotype of unstimulated CAR-T cells. *P* values in panels (**b**)-(**c**) were determined using a two-tailed unpaired student's t test (df = 4), with data shown being individual points along with means ± s.e.m.

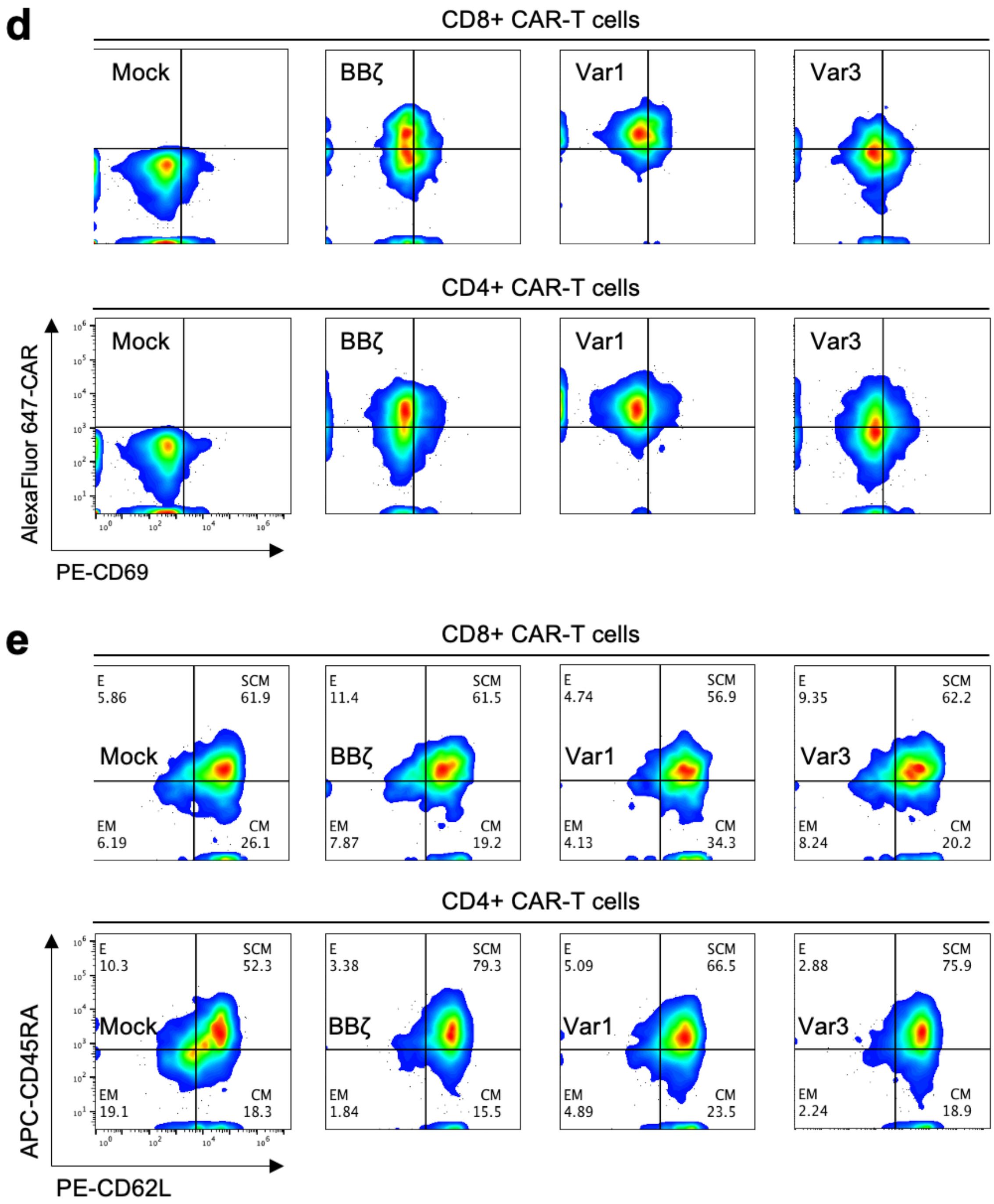


**Supplementary Figure 5 (continued).** Representations demonstrating basal phenotypic status of CAR-T cells in human primary CD4$^+$ and CD8$^+$ T cells. (**a**) FACS plots showing gating strategy. (**b**) Basal CD69 and (**c**) PD-1 expression in unstimulated human primary CD4$^+$ and CD8$^+$ CAR-T cells (n = 3 technical replicates representative of 2 biological replicates). *P* values for CD69 expression are 0.0050 for Var1 vs. BBζ and 0.0108 for Var3 vs. BBζ in CD8$^+$ T cells. *P* values for PD-1 expression are 0.0020 for Var3 vs. BBζ and 0.0020 for Var1 vs. Var3. **(d)** FACS plots showing CAR expression vs. CD69 expression in unstimulated CAR-T. (**e**) FACS plots showing memory phenotype of unstimulated CAR-T cells. *P* values in panels (**b**)-(**c**) were determined using a two-tailed unpaired student's t test (df = 4), with data shown being individual points along with means ± s.e.m.

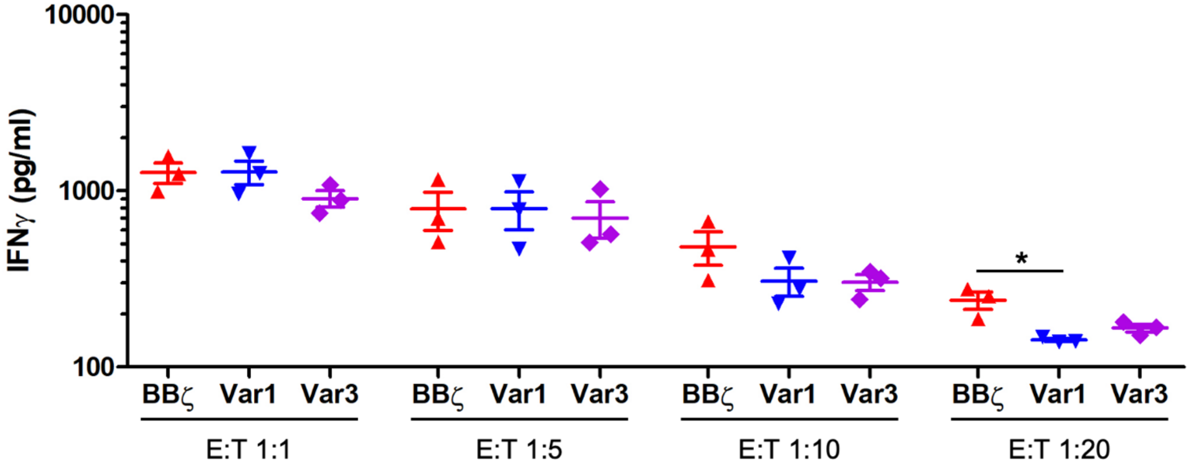


**Supplementary Figure 6.** IFNγ secretion levels of human primary CD8$^+$ CAR-T cells upon antigen challenge with NALM6 cells at various effector to target (E:T) cell ratios. The *p* value shown for Var1 vs. BB**ζ** at an E:T ratio of 1:20 is 0.0232 (df = 4). Data shown are individual points along with means ± s.e.m.

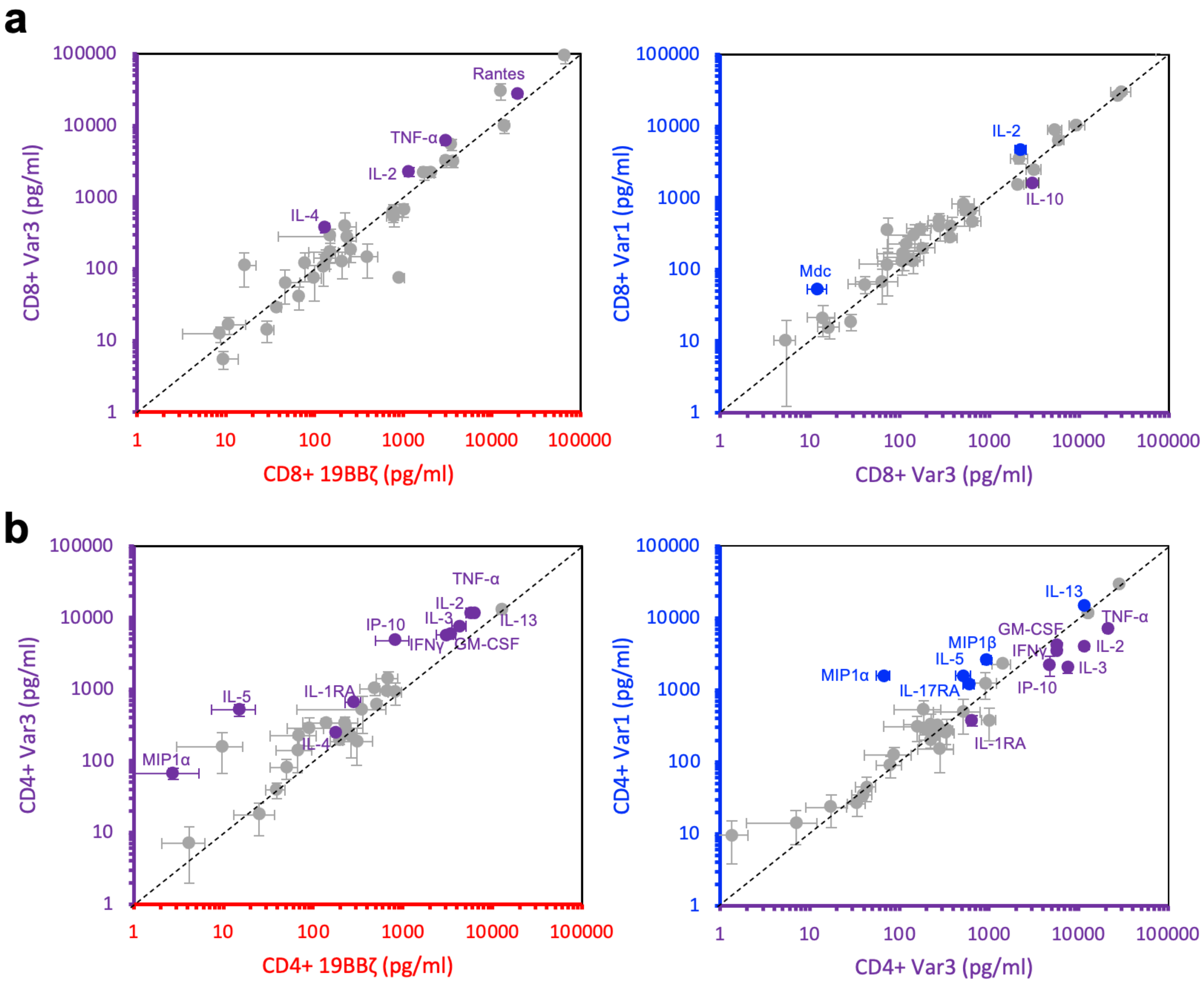


**Supplementary Figure 7.** Cytokine/chemokine profiles of human primary CD4+ and CD8+ CAR-T cells upon stimulation with NALM6 cells at an E:T ratio of 1:1.

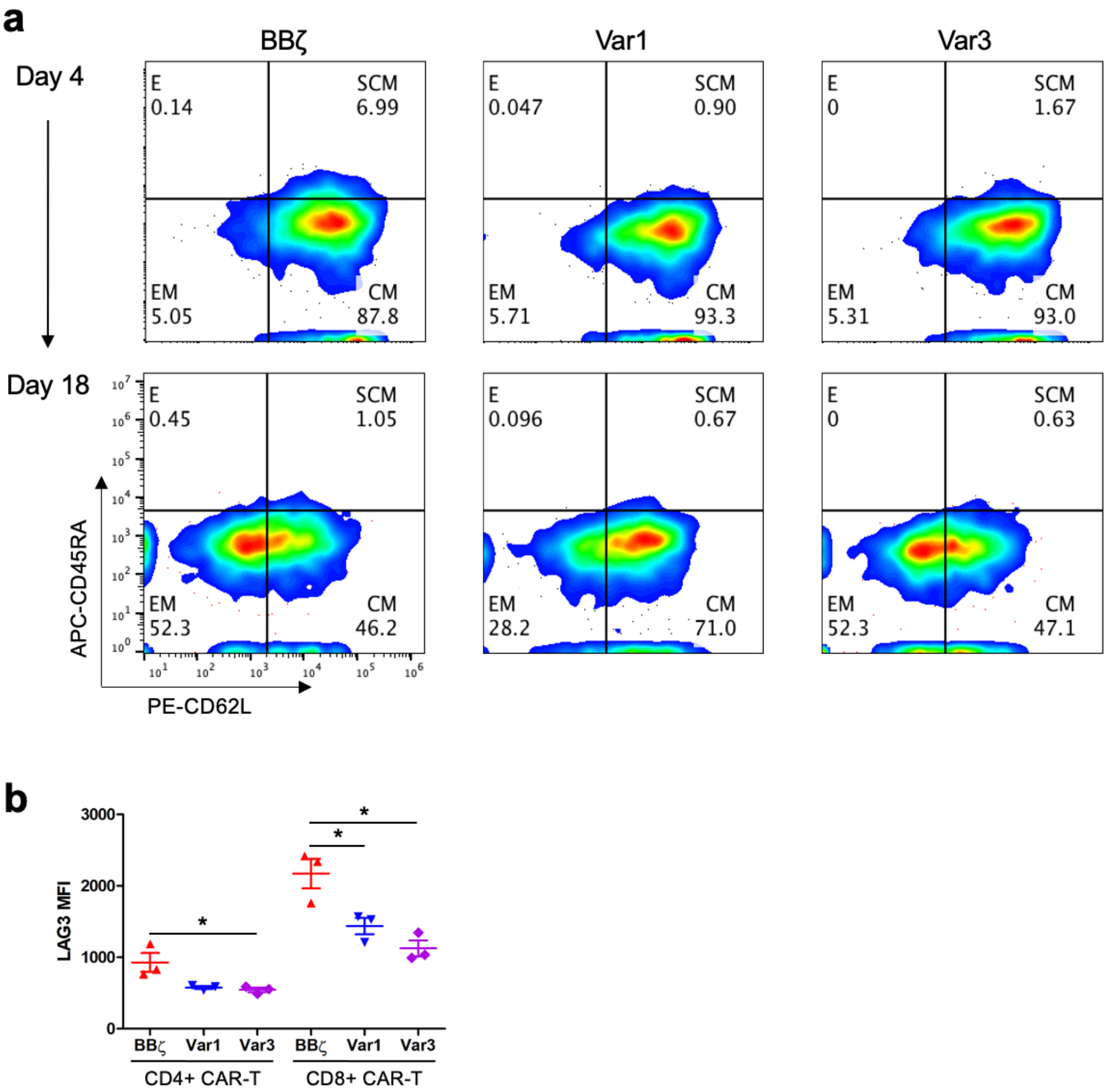


**Supplementary Figure 8.** (**a**) FACS plots displaying changes in CAR-T cell memory differentiation status following repeated tumor challenge (days 4 and 18 shown). (**b**) LAG-3 expression following repeated tumor challenge on day 22 (n = 3 technical replicates representative of 2 biological replicates). *P* values are 0.0465 for Var3 vs. BBζ in $CD4^+$ T cells, and 0.0358 for Var1 vs. BBζ and 0.0112 for Var3 vs. BBζ in $CD8^+$ T cell (df = 4). Data shown are means ± s.e.m.

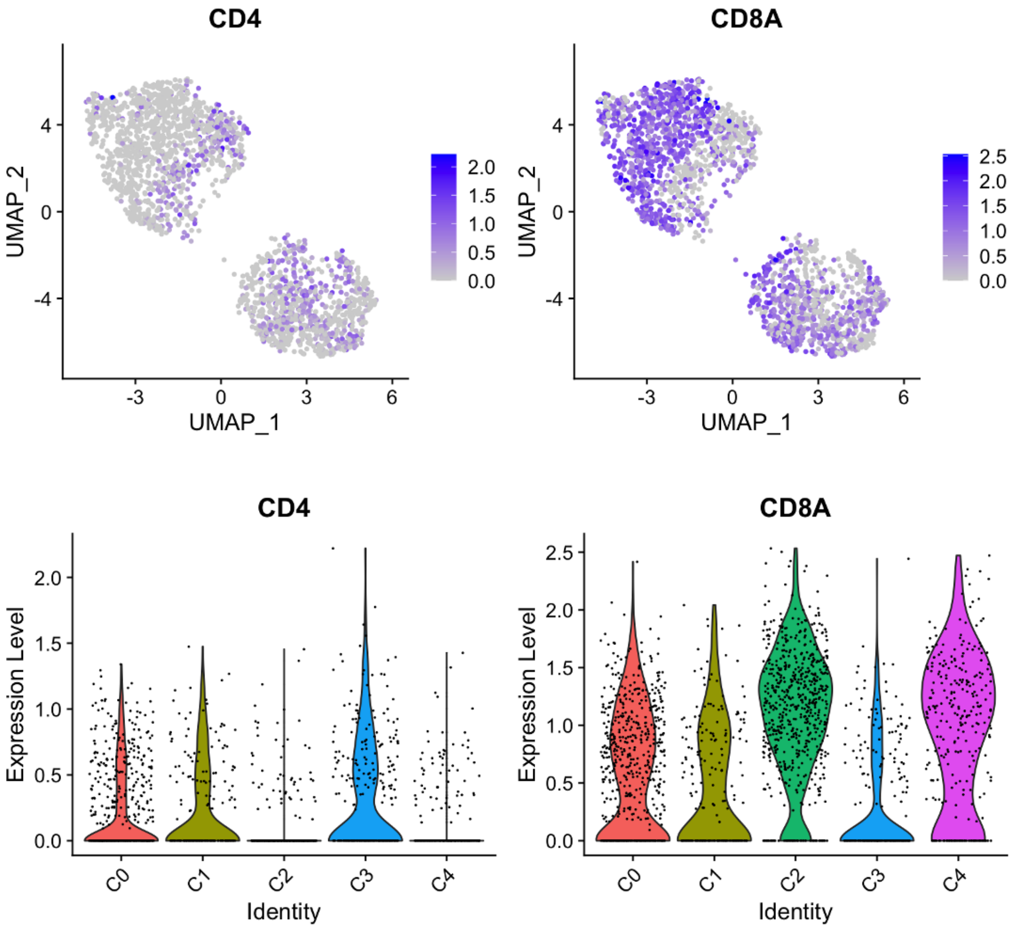


**Supplementary Figure 9.** Expression of *CD4* and *CD8A* across different phenotypic clusters. Related to **Fig. 3a.** Data points shown represent individual cells.

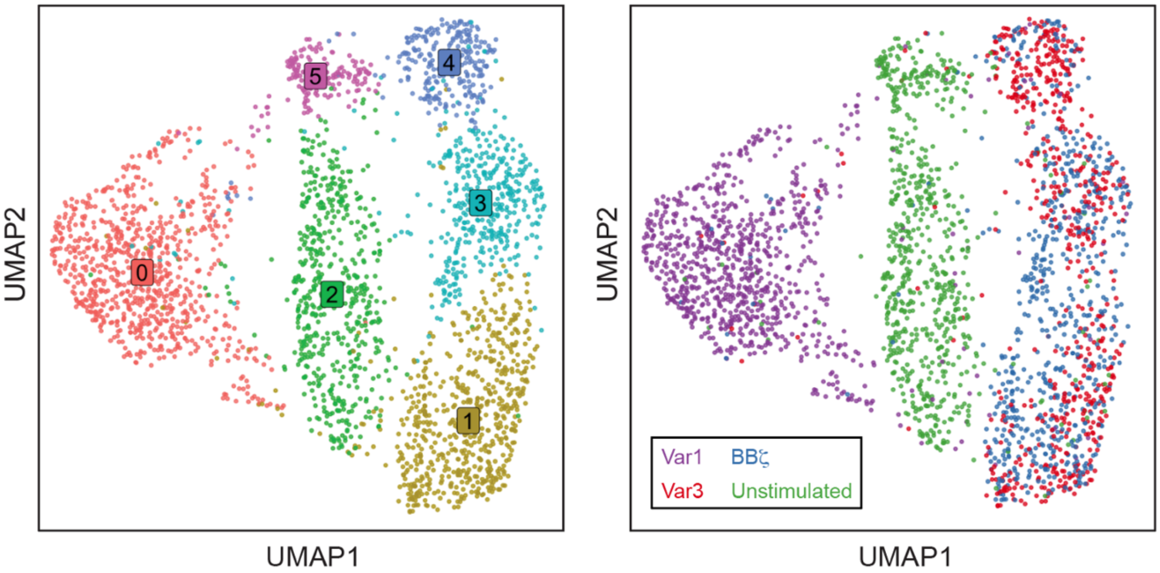


**Supplementary Figure 10.** Unsupervised clustering and visualization by uniform manifold approximation and projection (UMAP) of the scRNA-seq dataset, including 19BB**ζ**, Var1, and Var3 cells, as well as unstimulated, untransduced T cells (n = 3,086 total cells), colored by (**a**) cell state or by (**b**) sample. Data points shown represent individual cells.

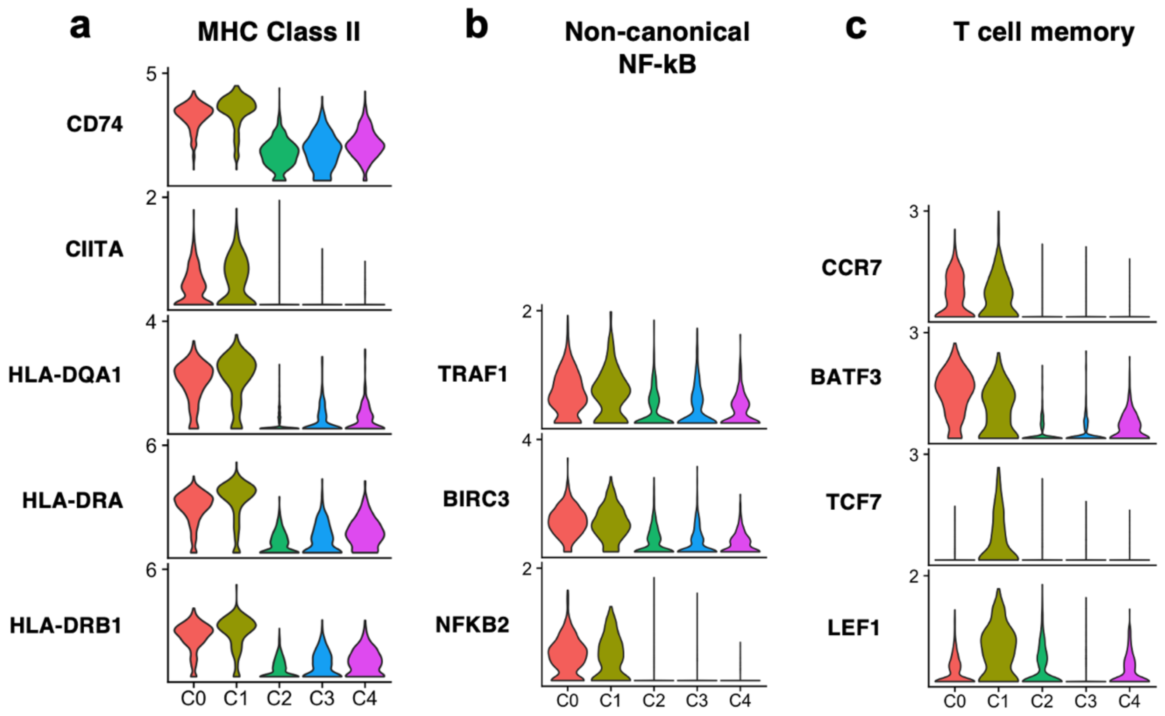


**Supplementary Figure 11.** Expression of genes related to (**a**) MHC class II, (**b**) non-canonical NF-kB signaling, and (**c**) T cell memory across cell state clusters.

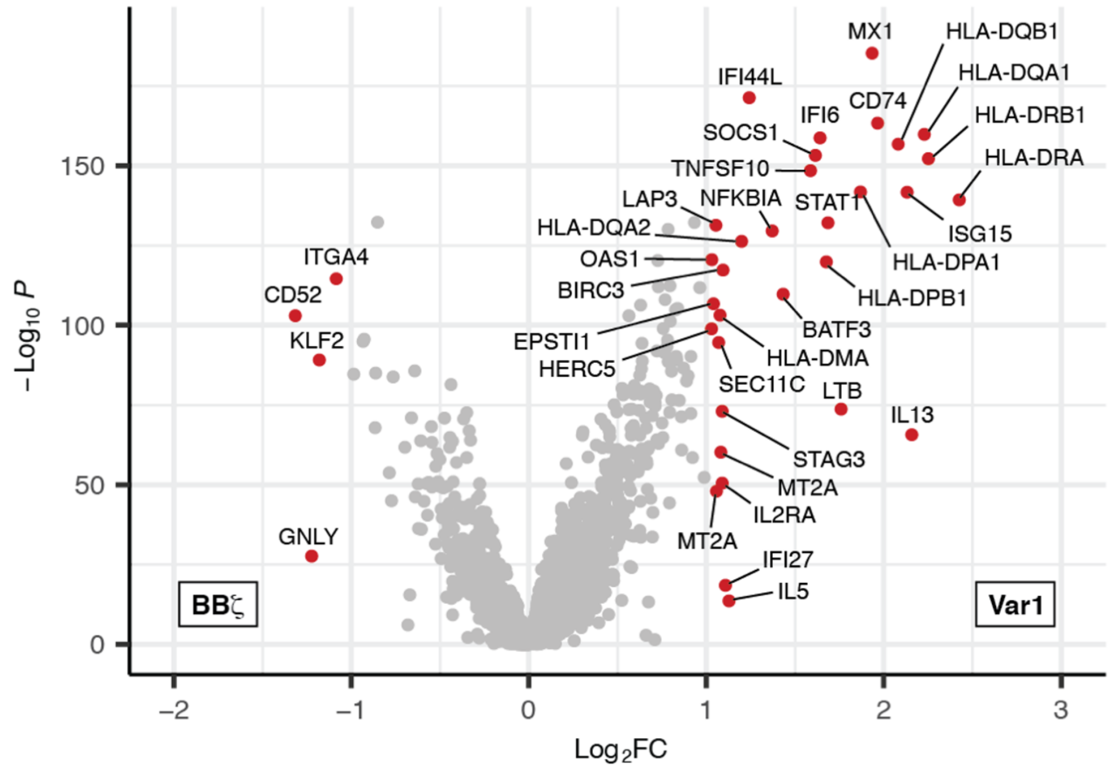


**Supplementary Figure 12.** Volcano plot showing differentially expressed genes between Var1 (n = 863 cells) and 19BB**ζ**-expressing (n = 637 cells) CAR T cells 48h after a third NALM6 rechallenge. *P* values for each gene were determined using a Wilcoxon Rank-Sum test. Genes with both a Bonferroni-corrected *P* value less than 0.05 and an average log2FC of > 1 are highlighted.

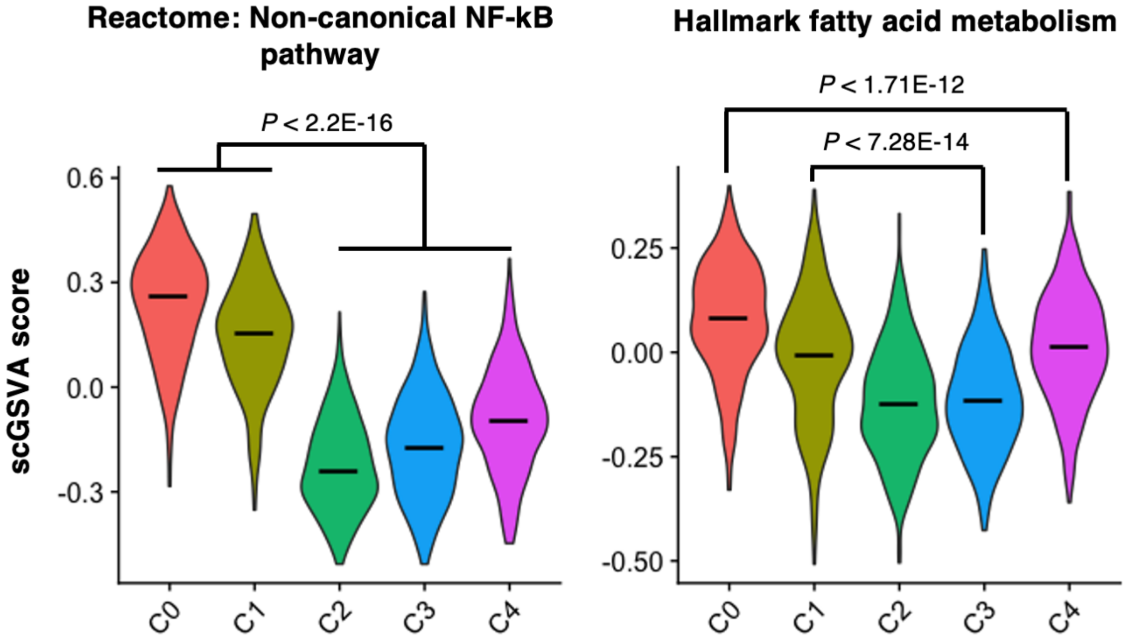


**Supplementary Figure 13.** Single-cell gene set variation analysis (scGSVA) scores from pathways conventionally associated with 19BBζ CARs across cell state clusters. Bars represent median scGSVA scores. *P* values were determined using a Wilcoxon Rank-Sum test.

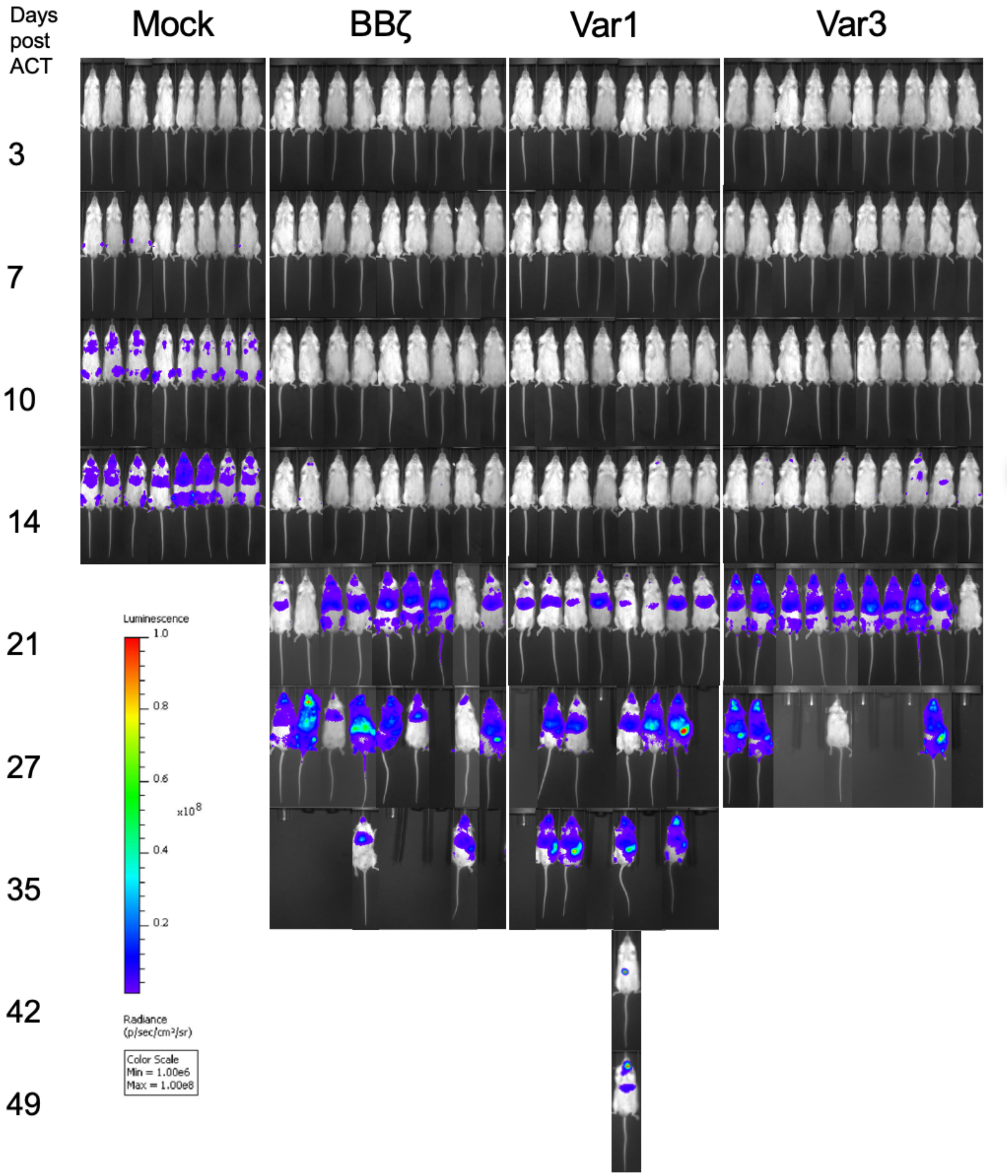


**Supplementary Figure 14.** Antileukemic activity of CAR-T cells *in vivo*. NOD-scid IL2ry$^{null}$ (NSG) mice were injected intravenously with FLuc+ CD19+ NALM6 cells, followed by treatment with 2x10$^{5}$ CAR-T cells i.v. 4 days later. Tumor burden was assessed periodically using an IVIS imaging system.

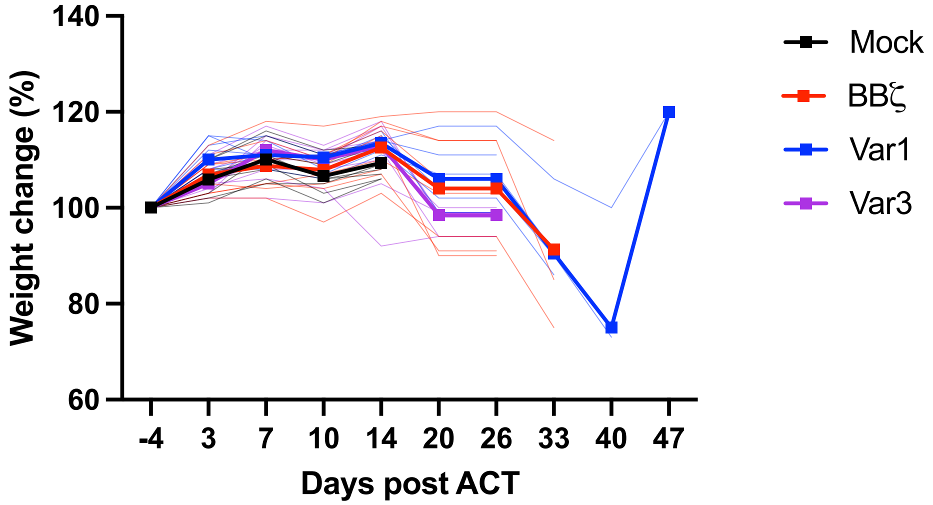


**Supplementary Figure 15.** Weight loss following treatment with $2x10^5$ CAR-T cells.

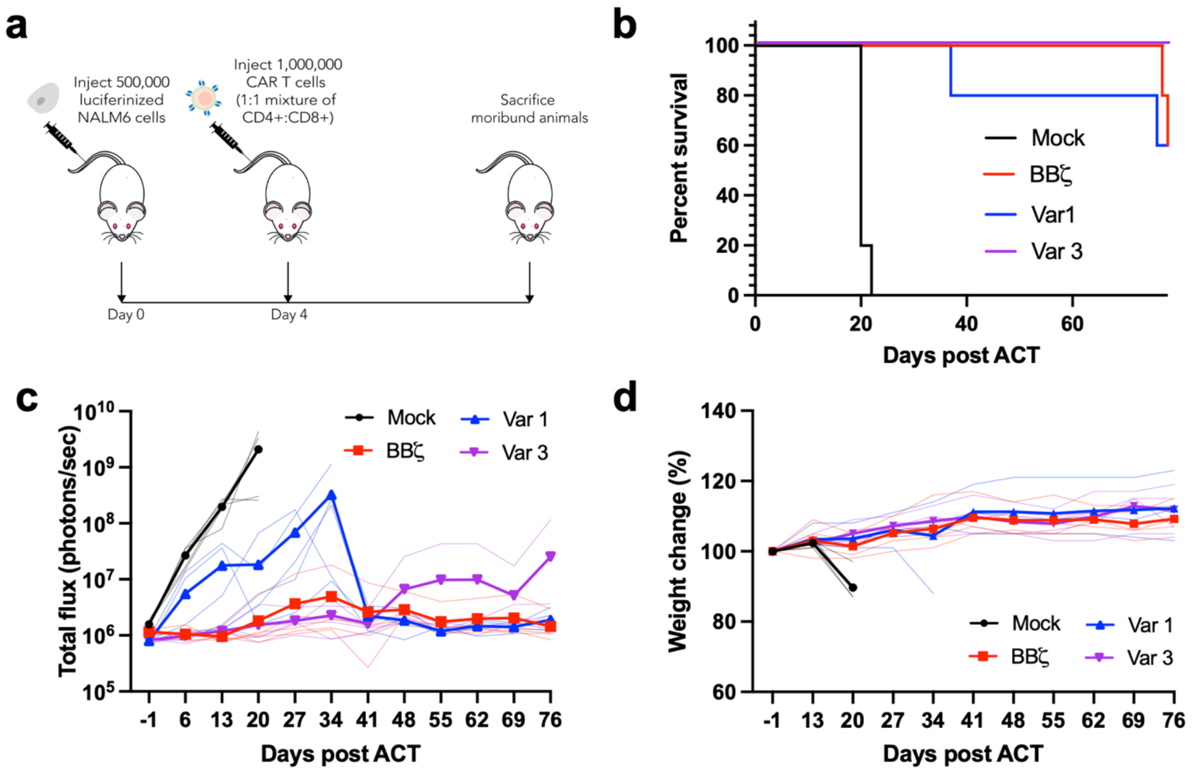


**Supplementary Figure 16.** High dose Var1, Var3, and BBζ achieve sustained remission. (**a**) Experimental design. NSG mice were infused intravenously with $5x10^5$ $FLuc^+$ $CD19^+$ NALM6 cells, then treated i.v. with $1x10^6$ of a 1:1 mixture of human $CD8^+$ and $CD4^+$ CAR T cells or untransduced control T cells (n = 5 mice for all groups). (**b**) Kaplan-Meier curve for overall survival. Tumor burden was assessed by measuring luminescent activity. (**c**) Quantification of total photon counts are shown. (**d**) Weight loss following ACT was monitored routinely. Data shown in panels (**c**)-(**d**) are individual points along with means shown in bold.

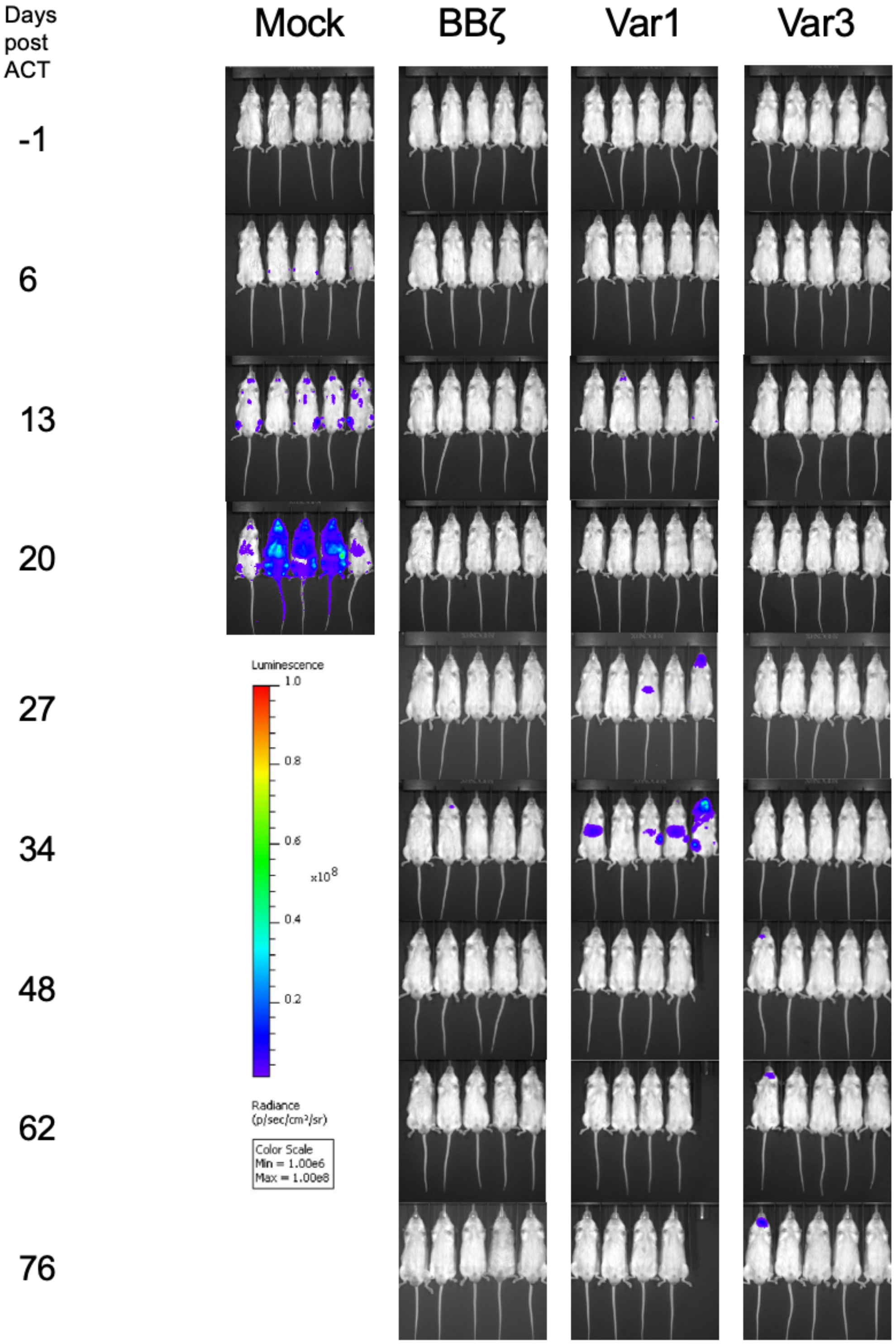


**Supplementary Figure 17.** Antileukemic activity of CAR-T cells *in vivo*. NOD-scid IL2ry$^{null}$ (NSG) mice were injected intravenously with FLuc$^{+}$ CD19$^{+}$ NALM6 cells, followed by treatment i.v. with 1x10$^{6}$ CAR-T cells 4 days later. Tumor burden was assessed periodically using an IVIS imaging system.
